# Enhanced perfusion following exposure to radiotherapy: a theoretical investigation

**DOI:** 10.1101/2023.06.12.544532

**Authors:** Jakub Köry, Vedang Narain, Bernadette J. Stolz, Jakob Kaeppler, Bostjan Markelc, Ruth J. Muschel, Philip K. Maini, Joe M. Pitt-Francis, Helen M. Byrne

**Author notes:** Equally contributing authors (Jakub Köry, Vedang Narain).

## Abstract

Tumour angiogenesis leads to the formation of blood vessels that are structurally and spatially heterogeneous. Poor blood perfusion, in conjunction with increased hypoxia and oxygen heterogeneity, impairs a tumour’s response to radiotherapy. The optimal strategy for enhancing tumour perfusion remains unclear, preventing its regular deployment in combination therapies. In this work, we first identify vascular architectural features that correlate with enhanced perfusion following radiotherapy, using *in vivo* imaging data from vascular tumours. Then, we present a novel computational model to determine the relationship between these architectural features, blood perfusion, and tumour response to radiotherapy *in silico*. If perfusion is defined to be the proportion of vessels that support blood flow, we find that vascular networks with small mean diameters and large numbers of angiogenic sprouts show the largest increases in perfusion post-irradiation for both biological and synthetic tumours. We also identify cases where perfusion increases due to the pruning of hypoperfused vessels, rather than blood being rerouted. These results indicate the importance of considering network composition when determining the optimal irradiation strategy. In the future, we aim to use our findings to identify tumours that are good candidates for perfusion enhancement and to improve the efficacy of combination therapies.

**Author summary:** Dysregulated tumour vasculature often contains hypoperfused blood vessels which inhibit the delivery of blood-borne anticancer therapies. Radiotherapy, used to treat more than half of all cancer patients, causes DNA damage to vascular endothelial cells, preferentially impacting smaller vessels, leading to their death and vessel pruning. At the same time, experiments measuring changes in tumour perfusion post-irradiation produce varying outcomes and, therefore, the impact of irradiation-induced vessel pruning on network-scale perfusion remains unclear. In this study, we use recent (*in vivo*) imaging data to identify features of tumour vascular architectures that impact perfusion change post-irradiation. We then use a newly-developed computational framework, directly informed by the experimental observations, to elucidate the relationship between the vascular geometry and topology prior to radiotherapy and the irradiation-induced changes to network perfusion. We find that perfusion increases most significantly for networks of blood vessels with small mean diameters and large numbers of angiogenic sprouts. Our results also distinguish different mechanisms of perfusion increase and we identify cases where rerouting of blood flow causes previously hypoperfused vessels to become perfused. Our study sheds more light on the impact of radiotherapy on tumour blood flow; these insights could be useful for improving anti-cancer treatments.

## 1 Introduction

The disordered architecture of tumour vasculature leads to poor perfusion, increased hypoxia, and oxygen heterogeneity. These factors impair a tumour’s response to oncological therapies [1].

The efficacy of radiotherapy, which is used to treat more than half of all cancer patients, depends on the generation of reactive oxygen species that induce irreparable DNA damage [1]. Pockets of hypoxia within a tumour can exhibit a lesser response to radiotherapy by up to a factor of three [2]. Moreover, adaptations to hypoxia can lead to tumour phenotypes with increased chemoresistance and metastatic capabilities [3]. For a list of drugs that perform more poorly in hypoxic conditions than in normoxic conditions, see [4].

Vessel normalisation (i.e. the pruning of redundant structures and remodelling of abnormal architectures) is generally considered favourable for oncological therapies [5]. In particular, pruning smaller, less mature vessels may help restore the balance between pro- and anti-angiogenic factors in the microenvironment [1, 6]. Therapies that can be used to accomplish this include radiotherapy, which preferentially impacts smaller vessels [7, 8]. Pruning of these vessels could alter the blood flow so that the ‘normalised’ vasculature is more conducive to the delivery of nutrients and therapeutics [6].

Therefore, the need to understand the normalisation of tumour vasculature is apparent. However, the biological mechanism behind this process is not fully understood. Indeed, experiments conducted to measure the change in tumour perfusion after irradiation of tumours have recorded varying outcomes.

A review by Park et al. found that reported results were inconsistent, and that functionality improved and then worsened in human tumour vasculature during the early stages of radiotherapy [9]. Conversely, in human tumour xenografts or murine tumours, irradiation resulted in mild to severe damage depending on the dosage, reducing blood perfusion [9].

Kim et al. found that stereotactic body radiation therapy decreased perfusion [10]. On the other hand, Shibuya et al. found that in the case of cervical cancer, blood flow improved significantly after irradiation [11]. Moreover, a study by Bussink et al. found that irradiation led to rapid changes in perfusion, observing increases shortly after irradiation followed by significant decreases [12].

Kaeppler et al. studied the response to single-dose and fractionated radiotherapy of two tumour cell lines — a highly vascular colon adenocarcinoma (MC38) and a less vascular melanoma (B16F10) — and found that the former contained a larger proportion of hypoperfused vessels compared to the latter [8]. Moreover, it was observed that the smaller-diameter vessels were more likely to be hypoperfused and also more likely to be pruned following irradiation (i.e. their endothelial cells were more susceptible to apoptosis). Thus, it was concluded that tumour perfusion post-irradiation depends on the density of small (typically hypoperfused) vessels.

An improved understanding of the relationship between irradiation and perfusion would allow for accurately-planned fractionated radiotherapy. However, it remains unclear why certain tumours exhibit increases versus decreases in perfusion after irradiation. The blood vessels that comprise tumour vasculature have heterogeneous architectural characteristics that present a complex system resistant to experimental analysis. Thus, there is a need for computational tools to guide and complement experimental research.

Broadly speaking, tumour vascular architecture has been modelled spatially with synthetic networks that reflect biological characteristics or, more recently, by digitising the morphology of real tumour vasculature.

Hierarchical and ordered vasculature has been modelled as a regular forking network by Bernabeu et al. using pathological dimensions gleaned from experimental data [13]. Non-hierarchical and ordered vasculature has been previously represented as a hexagonal network, similar to that observed in avian yolk sacs [14]. Alarcón et al. have used a similar network to couple processes at the intracellular, cellular, and tissue scales [15]. Owen et al. employed both hexagonal and disordered networks to show that vascular remodelling could be achieved with a balance of pruning and angiogenesis [16].

Synthetic networks are also commonly generated by simulating the random migration of capillary tips based on a chemotactic gradient. Anderson and Chaplain developed a model using a random walk biased towards higher TAF (tumour angiogenesis factor) levels, which resembled *in vivo* angiogenic networks [17]. Similar models of biased random motility and sprout formation have been employed by Macklin et al. and Shirinifard et al. in 2D and 3D, respectively [18, 19]. In addition, Stepanova et al. have employed a multiscale random walk model to study endothelial cell dynamics [20].

While synthetic networks can replicate many features of real tumour vasculature, some studies have gone a step further and digitised experimentally-acquired tumour vasculature. Grimes et al. used a 3D digitised network to estimate oxygen distribution [21]. Grogan et al. used a similar method to compare 2D and 3D representations, while Sweeney et al. used a 3D model to suggest that using realistic vasculature was key to modelling tumour fluid dynamics [22, 23].

Topological data analysis (TDA) is an emerging mathematical field that uses topological and geometric approaches to quantify the ‘shape’ of data [24, 25]. TDA characterises shape via topological invariants such as connected components and loops at multiple scales. Persistent homology, a method from TDA, has been successfully applied to quantify dynamic characteristics of vascular networks in experimental data [26, 27] and to distinguish between synthetic vascular networks produced by different parameter regimes in a mathematical model of tumour-induced angiogenesis [28]. In experimental data, the change in the number of loops in vascular networks following radiotherapy has shown great variation across different tumours relative to the day of irradiation (single dose) [27]. However, this study did not take perfusion into account and, in particular, did not differentiate between different initial structural characteristics of the networks pre-therapy which can affect their response to radiotherapy. More recently, a novel topological descriptor for weighted directed graphs was developed [29]. For a weighted directed graph, e.g. a vascular network weighted by vessel diameter or flow time with the directions given by the flow, the method outputs a barcode describing its structure. While this topological descriptor encodes rich information about such a network, it has not yet been applied to real data, is computationally intensive, and would need to be substantially compressed to enable interpretable comparisons of multiple networks. Motivated by previous TDA studies of tumour vasculature [26, 27, 28], we focus on the readily computable number of loops combined with geometric information. None of the previous studies directly explored the link between network topology and perfusion as altered following irradiation and to the best of our knowledge, there exists no experimentally-motivated computational study of the effect of irradiation-induced pruning on network perfusion that considers both geometric and topological architectural features.

In this paper, we aim to characterise the extent to which radiotherapy-induced changes in vascular morphology (e.g. pruning of small vessels) may contribute to increased vascular perfusion. While [8] provided a broad analysis comparing two tumour cell lines under various radiotherapeutic treatment regimes, we noticed that even within one cell line (MC38) under single-dose radiotherapy, some tumours increased and some decreased their perfusion. We will therefore focus solely on MC38 under single-dose radiotherapy and aim to assess whether certain structural features pre-irradiation can be used as a proxy for predicting perfusion response to radiotherapy.

The structure of this work is summarised in Figure 1. Having introduced the biological and computational background for this study, we describe the *in vivo* experimental study motivating and informing our model in Section 2. In this section, we also highlight key correlations between certain architectural (geometric and topological) metrics and changes in perfusion after irradiation. In Section 3, we introduce key aspects of our computational model including the proposed architecture of the initial networks, the pruning rules, and the way in which perfusion is measured. We then use our model to investigate the underlying causal links and present two mechanisms of perfusion improvement in Section 4. We contextualise our findings and discuss their implications in Section 5. The complete experimental procedure and model description can be found in Section 6.

**Figure 1:**
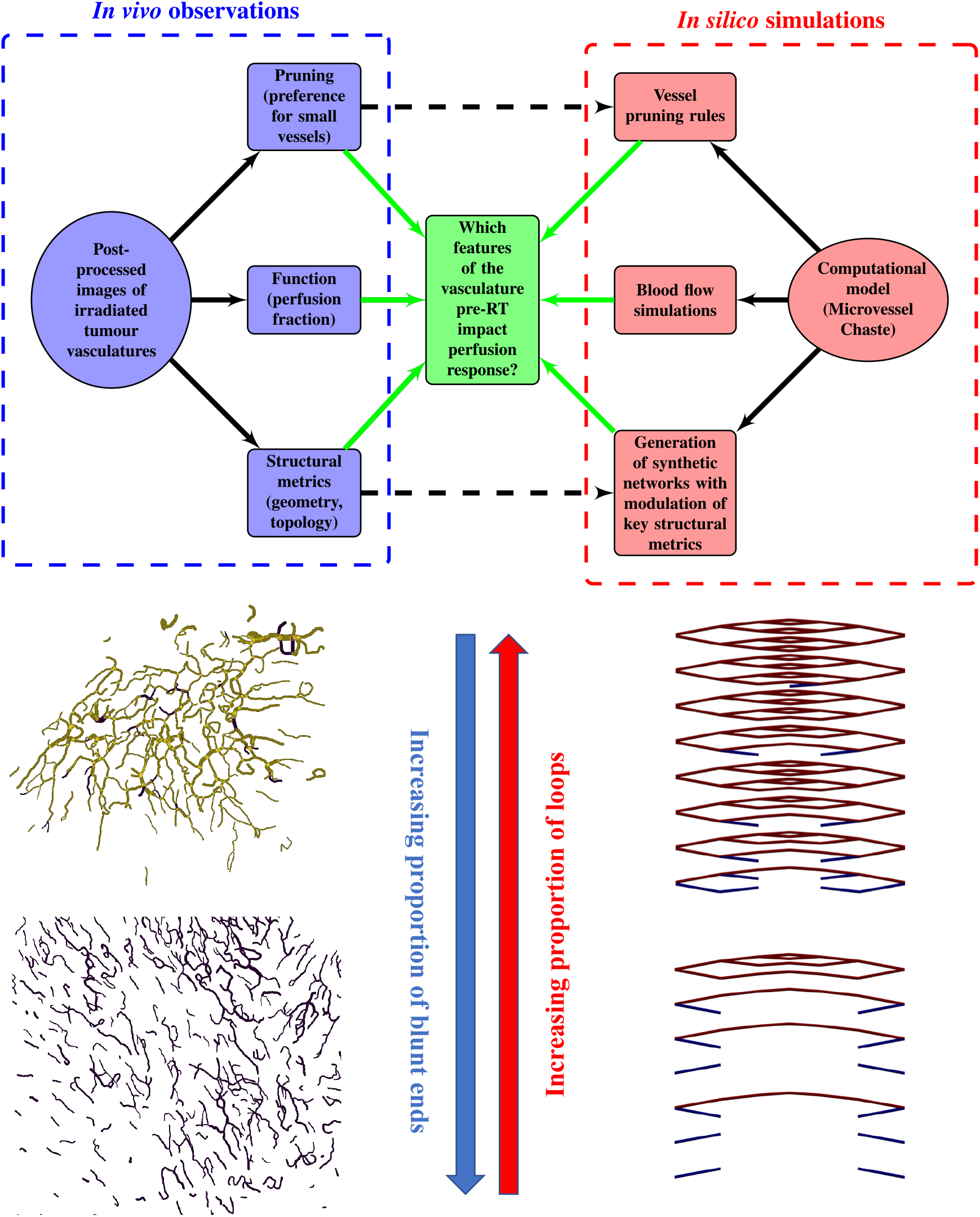
Chart summarizing the key components of the present study. Bottom-left panels present representative vascular regions containing perfused (yellow) and hypoperfused (purple) vessels from Day 0 (just before irradiation) for one of the tumours for which irradiation-induced vessel pruning led to a decrease in perfusion (tumour 6; top) and one for which it led to an increase (tumour 1; bottom). Note that the latter vasculature contained many hypoperfused blunt-ended vessels, while the former contained more loops. Bottom-right panels show pruned synthetic (forking) networks exhibiting similar properties with perfused (red) and hypoperfused (blue) vessels.

## 2 Experimental motivation

### 2.1 Uncertainty in direct flow simulations

Ideally, one would assess the impact of radiotherapy-induced vessel pruning on network perfusion by performing direct blood flow simulations on networks extracted from microscopy images. However, information about which network nodes serve as inlets and outlets is often limited or non-existent. Even given such information, it would still be challenging to determine whether constant pressure or constant flow rates should be imposed at inlets, and what values these pressures or flow rates should take to obtain quantitative agreement with measurements of blood flow *in vivo*. For these reasons, we choose a different strategy. First, in Section 2.2, we identify key global geometric and topological metrics that can be used to determine whether irradiation-induced vessel pruning improves or impairs the perfusion of vascular networks extracted from tumours. Then, in the following sections, we present a theoretical study of network perfusion and vessel pruning in synthetic networks. In doing so, we aim to confirm the utility of the newly-proposed metrics in distinguishing tumour vascular networks whose perfusion increases or decreases following radiotherapy.

### 2.2 Understanding perfusion response to radiotherapy and its determinants

Exposure to radiotherapy causes DNA damage in endothelial cells, which was found to induce cell death due to apoptosis preferentially in smaller hypoperfused vessels and result in cell cycle arrest in larger vessels which thus remain functional channels for the flow of blood [8, 30]. Regardless of the exact mechanism driving the cell death, we assume that radiation-induced DNA damage is the dominant cause of vessel pruning with a strong preference for small vessels and we neglect a preferential regression of hypoperfused vessels found in developmental vascular networks [31, 32].

We define perfused and hypoperfused vessels as in [8] (see also Section 6.1 for the experimental procedure and Section 3.3 for the minimum differentiating flow rate in our simulations) and similarly use the perfusion fraction (PF) to quantify network perfusion. The perfusion fraction (*t*) is the ratio of the number of perfused vessels to the total number of blood vessels, i.e.:

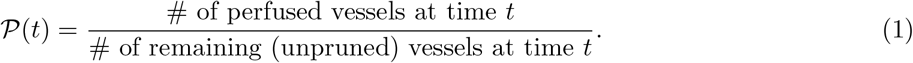

The perfusion fraction is time-dependent because both the total number of vessels and the number of perfused vessels change over time, due to pruning (see the representative example in Figure 2). Experimental observations of irradiated tumours suggest that vessel pruning occurs on timescales that range from hours to days [8, 30, 34]. We thus focus on short-term responses, during the first four days following radiotherapy (Figure 3). We number the studied tumours from 1 to 7.

**Figure 2:**
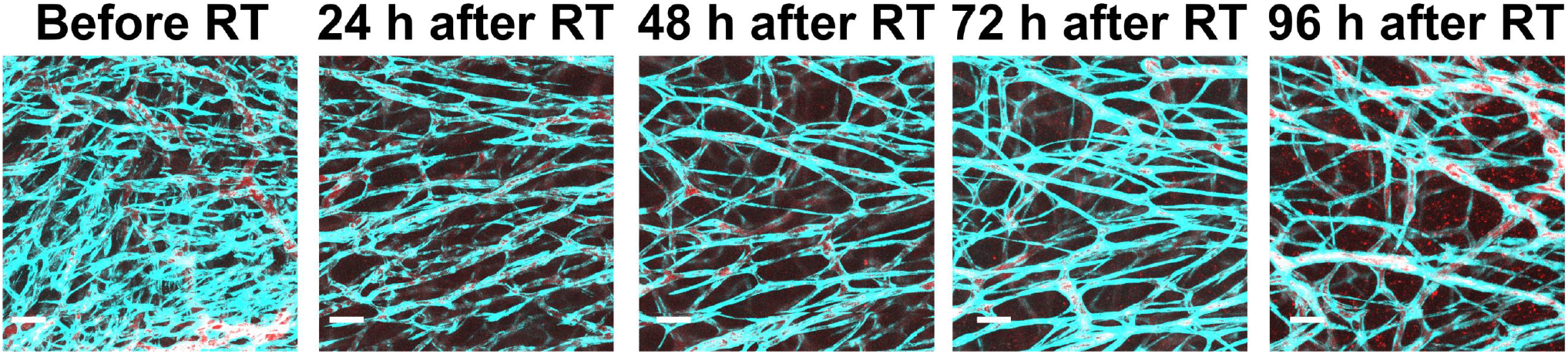
Representative vascular architectures from our experiments [8], where a single dose of 15 Gy X-rays was delivered to a tumour and changes to its vascular structure were monitored over the course of 4 days (reproduced from [33]). Endothelial cells are in cyan, while perfused vessels (qDot705) are in red. The scale bar corresponds to 100 μm.

**Figure 3:**
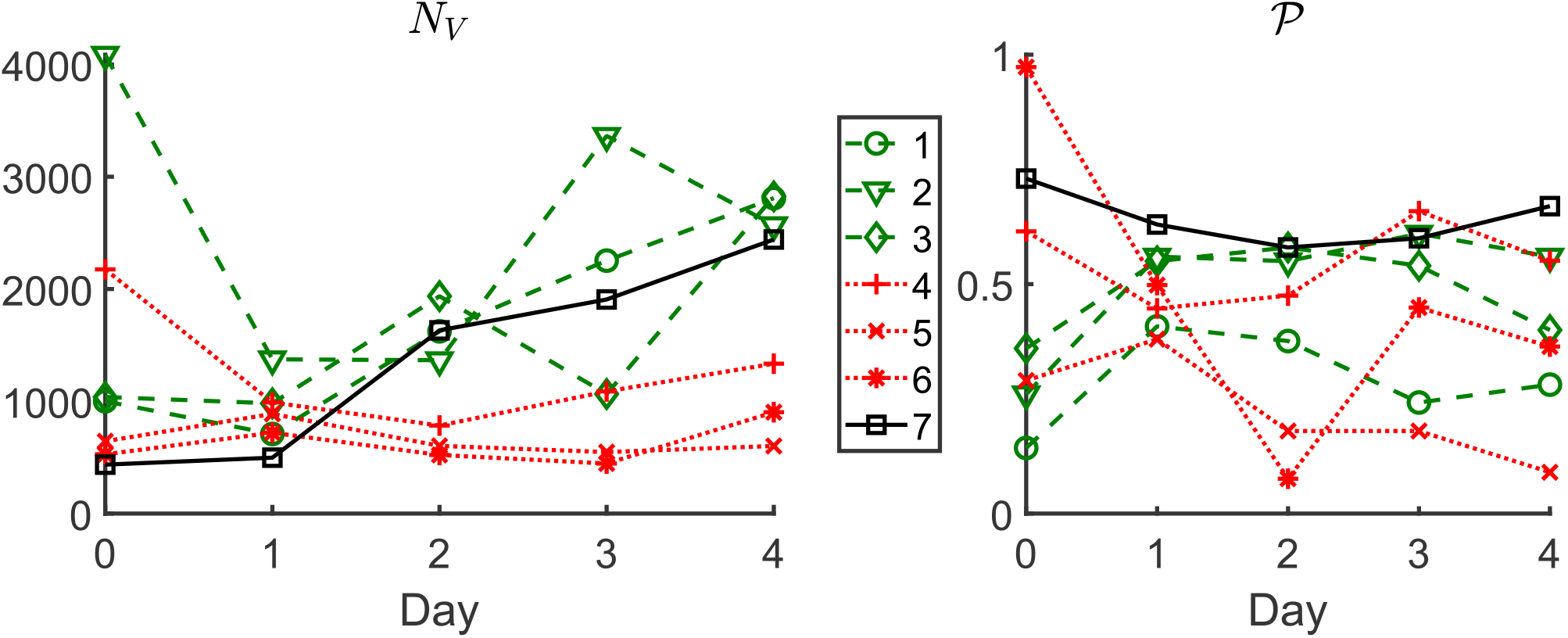
Time evolution over the first four days post radiotherapy (Day 0 is the day of irradiation) of vessel count *N*_*V*_ and perfusion fraction *𝒫* for the seven tumours studied in [8].

Firstly, we note that the time between irradiation (Day 0 in Figure 3) and the first day on which a reduction in vessel number (vessel pruning) was observed varied across the tumours that were studied. While most tumours (1, 2, 3, and 4) exhibited a decrease in vessel number on Day 1, others (5 and 6) experienced a slight increase in vessel number over this time period. We note further that this delayed decrease in vessel number was observed for tumours with a relatively low vessel number and (at the same time) a large average vessel diameter at the time of radiotherapy (see Table 2). We speculate that these tumours were still undergoing extensive angiogenesis and had not developed a fully functioning tumour microvasculature at the time of radiotherapy. We note also that the vessel count for tumour 7 increased up to, and including, Day 4. This tumour’s vasculature had the largest number of connected components per size, and the size of its largest connected component was the smallest across all tumours (see Table 5 in Appendix A). These data are indicative of problems in image processing for this tumour and, therefore, we exclude it from our analysis.

Secondly, the number of days over which the number of vessels decreased varied between tumours: for some tumours (1 and 3) the decrease only lasted one day, while for others (2, 4, 5, and 6) it lasted two days. This variability is likely due to the complex nature of the tumour microenvironment and uncertainties in the timescales for vessel pruning. The pruning phase was followed by a period characterised by a significant increase in vessel number, likely due to angiogenesis.

Based on the above observations, we divide the tumours into two groups A and B so that a tumour belongs to group A or B if its perfusion fraction increases or decreases respectively during the pruning phase, which we define to be the period from Day 0 to the last day on which the number of vessels decreased, prior to the onset of angiogenesis. For a more quantitative comparison, we also define the pruning-induced perfusion difference Δ *𝒫* and its relative counterpart Δ_%_ *𝒫* as follows:

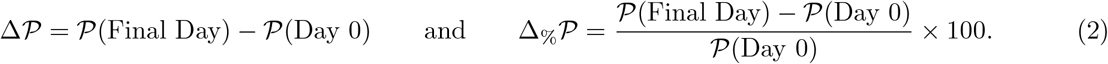

The perfusion increases if Δ_%_ *𝒫 >* 0 and decreases if Δ_%_ *𝒫 <* 0. In Table 1, we summarize these quantities and rank the tumours with respect to the relative change in PF (as measured by Δ_%_ *𝒫*).

**Table 1:** Perfusion fractions on Day 0 and the final day of the pruning phase, the pruning-induced perfusion difference, and its relative counterpart for six studied tumours. Tumours are ordered based on the relative change in PF as measured by Δ_%_ *𝒫*.

| Tumour | $\mathcal{P}(\text{Day 0})$ | $\mathcal{P}(\text{Final Day})$ | $\Delta\mathcal{P}$ | $\Delta\% \mathcal{P}$ |
| --- | --- | --- | --- | --- |
| 1 | 0.14 | 0.41 | 0.27 | 192.86 |
| 2 | 0.26 | 0.56 | 0.30 | 115.38 |
| 3 | 0.36 | 0.55 | 0.19 | 52.78 |
| 4 | 0.62 | 0.48 | -0.14 | -22.58 |
| 5 | 0.29 | 0.18 | -0.11 | -37.93 |
| 6 | 0.97 | 0.45 | -0.52 | -53.61 |

It might seem that the response to radiotherapy can be predicted by whether or not the tumour is initially highly-perfused. Tumours that are initially highly-perfused (e.g. 4 and 6) tend to have impaired perfusion post radiotherapy while those that are initially hypoperfused tend to have improved perfusion post radiotherapy (e.g. 1 and 2). However, even in our small dataset, we can find a counterexample: tumour 5 experienced a low perfusion fraction on Day 0, and an even lower value post-pruning. The particular value of the perfusion fraction (on any day of the experiment) is expected to be strongly dependent on the number, location, and strength of the inlet vessels which are often unknown. However, in the absence of such information and provided the inlet vessels are not pruned, it may still be possible to predict the perfusion response of tumour vasculature to radiotherapy (as measured by the relative change in the perfusion fraction Δ_%_ 𝒫) based on geometric and topological characteristics of the vasculature on Day 0 (i.e. just before the irradiation). Next, we carefully investigate the relationship between key characteristics of the tumour vasculature on Day 0 and the perfusion response.

#### 2.2.1 Network size (vessel count)

In general, the resistance of a vascular network to flow increases with the number of vessels. Therefore, one might naively expect that pruning vessels in a large network should always increase its perfusion and, consequently, that irradiation should improve perfusion at a rate proportional to the number of vessels in the network. Most of our networks are consistent with this principle. However, the PF decreased following irradiation for tumour 4, even though it has a large network (see Table 2). We conclude that the vessel count alone is insufficient to predict perfusion response to radiotherapy.

#### 2.2.2 Geometric determinants of perfusion

We now consider geometric determinants of flow resistance in individual vessels. One typically assumes that the flow rate *Q* through a cylindrical vessel, subject to a pressure drop Δ*p* across its length *L*, satisfies Poiseuille’s law:

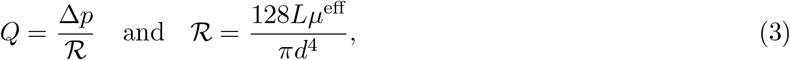

where *d* denotes vessel diameter and *μ*^eff^ is the effective viscosity of blood [13, 35]. This parameter depends in a nonlinear way on the vessel diameter and haematocrit (see [36]). Determining the haematocrit distribution within a network is challenging due to uncertainty in the locations and strengths of the inlets, the lack of consensus about the functional form describing haematocrit splitting (this remains an active research area [13, 37, 38, 39]), and the highly coupled and nonlinear nature of the haematocrit and blood flow. Therefore, for the sake of computational efficiency, we impose a uniform haematocrit *H* = 0.45 in all networks. Substituting this value into standard formulas obtained in [36] (see Equations (17)–(19)), we arrive at a relationship describing how the effective blood viscosity depends on vessel diameter. Note that for consistency with existing mathematical models [36], all length scales in Equations (17)–(19) are nondimensionalised with respect to 1 μm. Substituting this relationship into Equation (3), we obtain an explicit expression for the resistance ℛ in a vessel of length *L* and diameter *d*. As a further simplification, we propose the quotient ℛ ^*geom*^ = *L/d*^4^ as the simplest proxy for vessel resistance involving only the key geometric parameters. At the network scale, we define the following measures for the total resistance of a network:

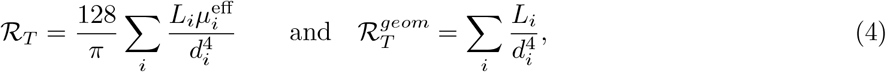

where the sum is over all network vessels *i*. Finally, we define the following proxies for mean vessel resistance:

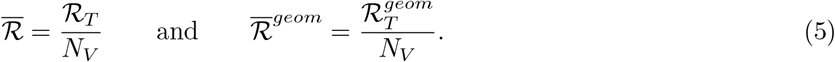

Due to the fourth-power dependence of the resistance on diameter, we expect that the average vessel diameter in a network will strongly impact its resistance. Table 2 confirms this: networks with small mean diameters typically have high mean resistance (both 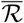 and 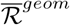). Furthermore, the mean diameter and mean resistances appear to be good predictors of radiotherapy response: low values of mean diameter (high values of mean resistance) correspond to group A, and vice versa. We note that neither the mean length nor the total resistance (which neglects network size, and decreases when vessels are pruned) distinguish between the two groups of tumours. In summary, for the six studied tumours, the mean diameter 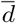 and mean resistance 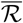 distinguish between the two groups, with group A having, on average, thinner (higher resistance) vessels than group B.

**Table 2:**
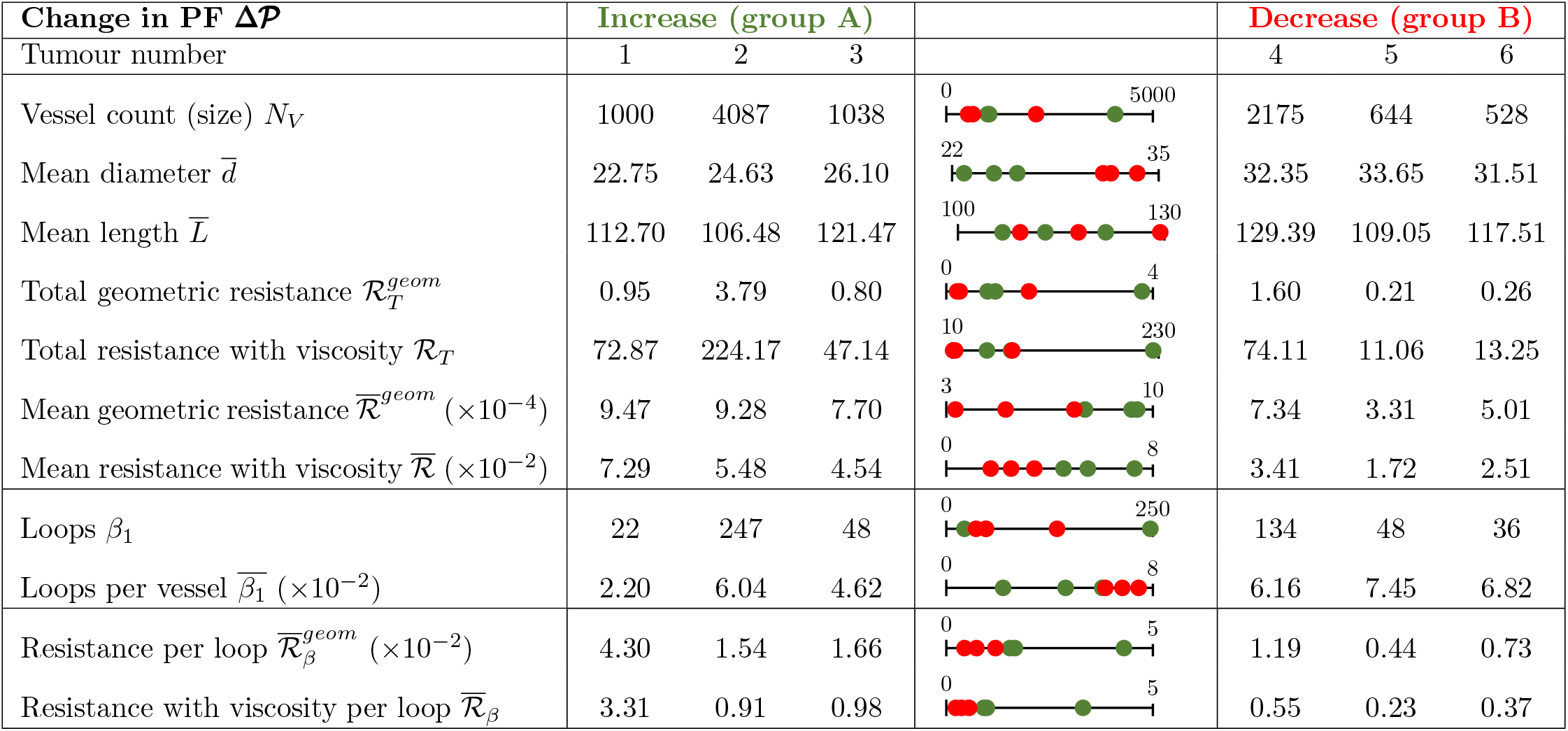
Key geometric and topological characteristics of tumour vasculatures on Day 0. Note that all lengths are in μm, all geometric resistances (total, mean, and per loop) in μm^*−*3^, and all resistances with viscosity in cP·μm^*−*3^, where cP (centipoise) is a unit of dynamic viscosity equal to mPa*s*).

| Change in PF $\Delta\mathcal{P}$ | Increase (group A) | | | | Decrease (group B) | | |
| --- | --- | --- | --- | --- | --- | --- | --- |
| Tumour number | 1 | 2 | 3 |  | 4 | 5 | 6 |
| Vessel count (size) $N_V$ | 1000 | 4087 | 1038 | | 2175 | 644 | 528 |
| Mean diameter $\bar{d}$ | 22.75 | 24.63 | 26.10 | | 32.35 | 33.65 | 31.51 |
| Mean length $\bar{L}$ | 112.70 | 106.48 | 121.47 | | 129.39 | 109.05 | 117.51 |
| Total geometric resistance $\mathcal{R}_T^{geom}$ | 0.95 | 3.79 | 0.80 | | 1.60 | 0.21 | 0.26 |
| Total resistance with viscosity $\mathcal{R}_T$ | 72.87 | 224.17 | 47.14 | | 74.11 | 11.06 | 13.25 |
| Mean geometric resistance $\bar{\mathcal{R}}^{geom} (\times 10^{-4})$ | 9.47 | 9.28 | 7.70 | | 7.34 | 3.31 | 5.01 |
| Mean resistance with viscosity $\bar{\mathcal{R}} (\times 10^{-2})$ | 7.29 | 5.48 | 4.54 | | 3.41 | 1.72 | 2.51 |
| Loops $\beta_1$ | 22 | 247 | 48 | | 134 | 48 | 36 |
| Loops per vessel $\bar{\beta}_1 (\times 10^{-2})$ | 2.20 | 6.04 | 4.62 | | 6.16 | 7.45 | 6.82 |
| Resistance per loop $\bar{\mathcal{R}}_\beta^{geom} (\times 10^{-2})$ | 4.30 | 1.54 | 1.66 | | 1.19 | 0.44 | 0.73 |
| Resistance with viscosity per loop $\bar{\mathcal{R}}_\beta$ | 3.31 | 0.91 | 0.98 | | 0.55 | 0.23 | 0.37 |

#### 2.2.3 Topological determinants of perfusion

We use Betti curves [40, 41] to track the number of loops in vascular networks during radiotherapy-induced vessel pruning. The Betti numbers *β*_0_ and *β*_1_ refer to the number of connected components and the number of loops in a network, respectively. Via the Euler-Poincaré formula [41, 42], the Euler characteristic *𝒳* of a network, which is given by its number of edges (vessels) *N*_*V*_ and nodes *N*_*N*_, i.e. *𝒳* = *N*_*N*_ *− N*_*V*_, is directly connected to its Betti numbers: *X* = *β*_0_ *− β*_1_. Given *β*_0_, *N*_*V*_, and *N*_*N*_, *β*_1_ can therefore be computed as [41]:

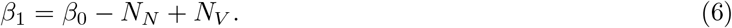

To obtain a Betti curve for *β*_1_, we track the number of loops throughout the pruning process on a network. As in the previous section, we also introduce metrics normalised by the number of vessels, i.e.:

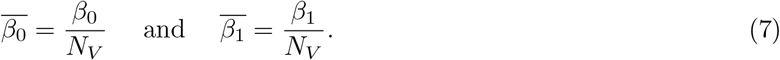

We note from Tables 2 and 5 that *β*_0_, 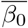 and *β*_1_ do not distinguish between the two groups of tumours (A and B), whereas 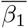 — which we will call loops per vessel — provides a (weak) distinction between the two groups: tumours with fewer loops per vessel belong to group A. To elucidate why the loops per vessel on Day 0 might impact the perfusion response, we use Equations (6) and (7) to write:

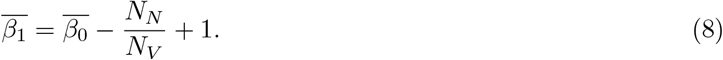

In what follows, we consider a network containing *N*_*I/O/ST*_ nodes of degree 1 (inlets, outlets, and tips of angiogenic sprouts that have not yet anastomosed) and *N*_*B*_ nodes of degree 3 (bifurcation points). For simplicity, we ignore nodes of degree 2 (vessels subdivided into segments), 4 (trifurcations), and higher as these seldom appear in our networks. We then have *N*_*N*_ = *N*_*I/O/ST*_ + *N*_*B*_ and, counting vessels twice by looping over all network nodes, we have *N*_*V*_ = (*N*_*I/O/ST*_ + 3*N*_*B*_)*/*2. Using these relations, Equation (8) can be simplified to read:

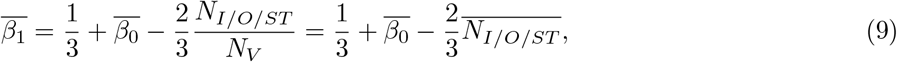

where 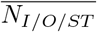 denotes the normalised number of degree-1 nodes. We see that small values of 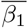 result from large numbers of nodes of degree 1, and vice-versa. As the degree of a node relates to its connectivity, vessels adjacent to degree-1 nodes are unlikely to play a significant role in network perfusion and pruning such vessels is desirable. Moreover, while some of these are inlet or outlet nodes, the majority are likely to represent tips of angiogenic sprouts (blunt ends) that have not yet anastomosed. Consequently, the relevant vessels are hypoperfused. Pruning these sprouts increases the perfusion fraction simply by reducing the denominator in Equation (1), whereas pruning vessels that connect bifurcation points (degree = 3) may disconnect parts of the network that were previously connected. This is confirmed in the bottom-left panels in Figure 1: the vasculature for which the PF increased (tumour 1; bottom) contains more angiogenic sprouts (i.e. is less inter-connected, with lower 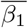) on Day 0, than the one for which the PF decreased (tumour 6; top). Taken together, these results provide a possible explanation for why tumours with low 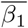 increase their PF, and vice versa.

#### 2.2.4 Metrics combining geometry and topology

A full understanding of the determinants of network perfusion is still lacking. It requires more detailed knowledge of the diameter and length distributions within a network and their connectivity. The intricate (nonlinear) nature of blood rheology further complicates the situation. A single measure (geometric or topological) is unlikely to contain complete information about the perfusion fraction. To assess the impact of radiotherapy on the perfusion fraction, one needs to consider the distribution of high- and low-resistance loops that are being pruned from the network. To this end, and inspired by the observations above, we now propose new metrics that combine network topology with its geometric properties in the form:

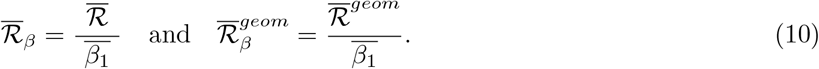

Table 2 shows that these metrics can distinguish between the two groups of tumours (particularly 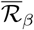 which accounts for the dependence of viscosity on the vessel diameter). To our knowledge, these are the first global metrics to combine the geometry and topology of vascular networks.

Taken together, our analysis suggests that networks containing large numbers of high-resistance vessels and angiogenic sprouts are likely to increase their PF following radiotherapy. In other words, we postulate that irradiation-induced vessel pruning will increase the PF of vascular networks containing large proportions of thin and blunt-ended vessels — we will next test this hypothesis using synthetic networks.

## 3. Model overview

Our analysis of the experiments revealed that vascular architecture influenced the outcome of pruning: tumours with lower mean vessel diameters 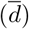 and fewer vascular loops per vessel 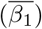 exhibited an increase in perfusion (Δ_%_𝒫 *>* 0) when irradiated. We then used our computational model to better understand the causal relationship between the architectural properties of a vascular network and how its perfusion changes following radiotherapy (Figure 4).

**Figure 4:**
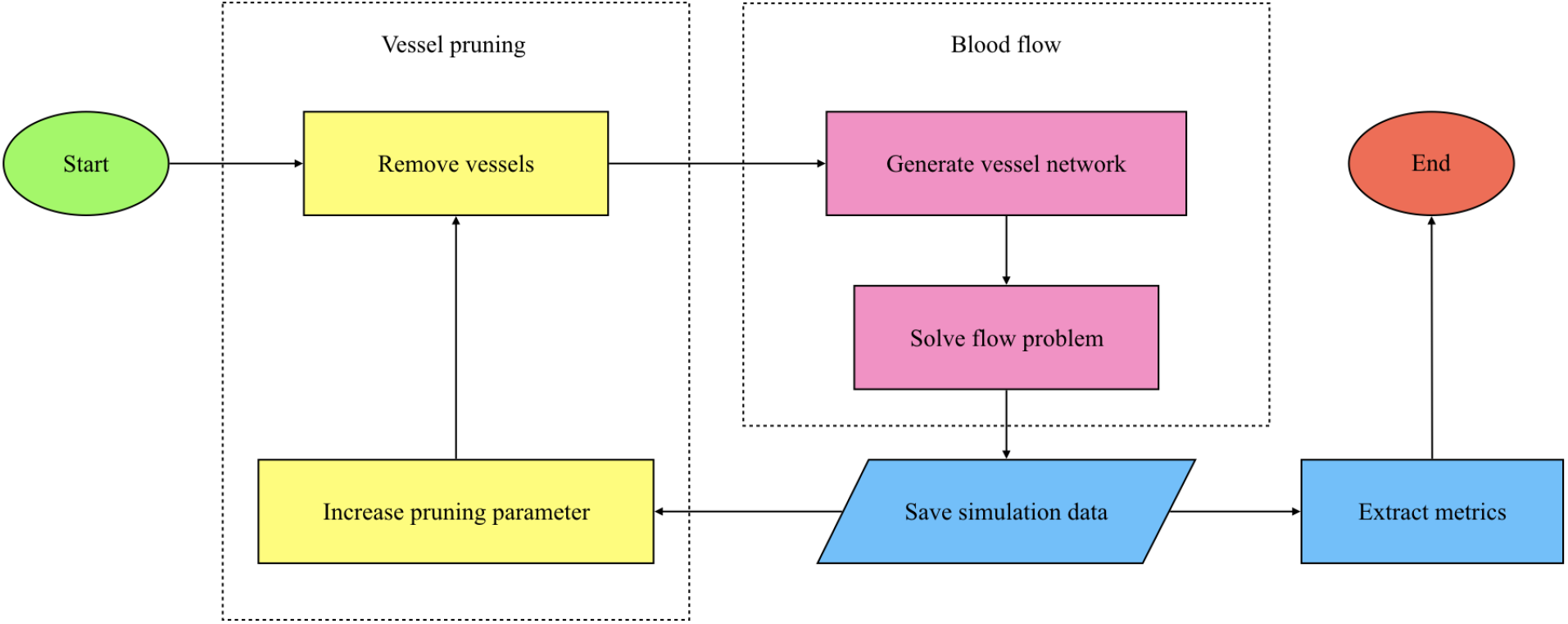
Flow chart summarising our model components. Simulations were carried out using Microvessel Chaste [43, 44].

In brief, we constructed multiple networks to replicate the characteristics of interest (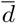 and 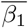) exhibited by the biological dataset. We then simulated blood flow through the networks and measured the change in perfusion as vessels were successively pruned. The design of the model is detailed below, with the parameters in Appendix E.

### 3.1 Network design

We used forking and hexagonal networks to represent hierarchical and non-hierarchical vasculature respectively (Table 3). We then simulated blood flow through the networks. Using simple symmetric geometries allows us to isolate the influence of the different geometric and topological factors (e.g. vessel diameters, lengths, and network loops) under consideration. While the geometries of the networks are simple, they reflect several properties of biological vasculature. Here, we describe the forking networks since they form the focus of our results. The design of the hexagonal network can be found in Appendix C.

**Table 3:** Heterogeneity in both hierarchical and non-hierarchical networks is modulated by a single parameter.

| Architecture | Hierarchy | Heterogeneity Parameter | Values |
| --- | --- | --- | --- |
| forking | present | relative thickness of daughter vessels ($\alpha$) | 1.1, 1.2, 1.3 |
| hexagonal | absent | SD of diameter distribution ($\sigma$) | 8.68 $\mu\text{m}$, 13.23 $\mu\text{m}$, 17.49 $\mu\text{m}$ |

In the forking network, each vessel divides into two daughter vessels until seven generations are created (counting the inlet vessel as generation 0). After the seventh generation (generation 6), the network converges symmetrically into a single outlet vessel. Note that throughout this work, for simplicity, we consider any straight-line segment to be an individual vessel (as opposed to defining vessels by their two endpoints being bifurcation nodes, inlets, outlets, or blunt ends).

#### 3.1.1 Vessel dimensions

The diameters of the two daughter vessels (*d*_*A*_ and *d*_*B*_) are related to the diameter of the parent vessel (*d*_*parent*_) via Murray’s Law [45]:

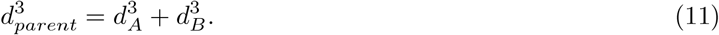

Therefore, the inlet vessel diameter *d*_*inlet*_ modulates the thickness of all subsequent vessels. Following [13], we assume that the vessel lengths *L* are proportional to vessel diameters (*d*) so that:

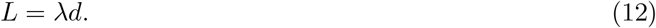

Based on the mean vessel length to mean diameter ratios in our experimental data, we fix *λ* = 4. The branching angle at a bifurcation is dictated by the *y*-extent of its constituent vessels. The *y*-extent of a vessel in generation *i≥* 1 is the length of its projection onto the *y*-axis (*V* _i_) in a manner that places the following limit on the spatial extent of the network [13]:

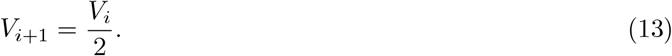

We set *V*_1_ = 0.9*L*_1_ for our simulations to allow the network to extend to a sufficient degree along the *y*-axis, in line with [13].

If we further impose that two daughter vessels of any parent vessel must have the same diameter (*d*_*A*_=*d*_*B*_), then the network geometry is fully specified. This forms our reference network (Figure 5).

**Figure 5:**
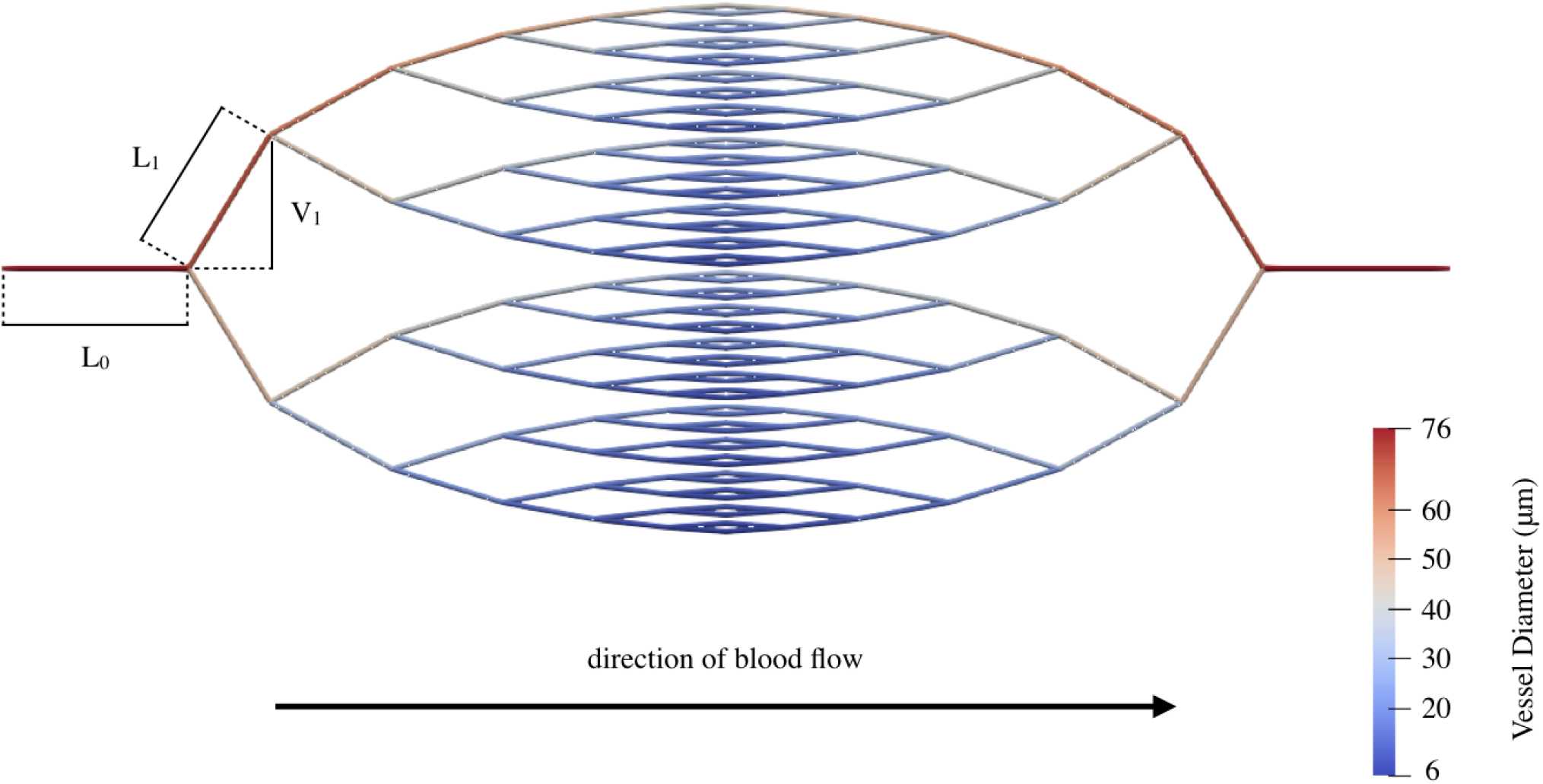
Blood flows from left to right through a single inlet and a single outlet in the forking network. *L*_i_ and *V*_i_ denote the length and *y*-extent of a vessel in generation *i*, respectively.

Our reference network must be modified to reflect the properties of interest (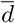 and 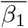) in the biological networks. Therefore, we offset the mean vessel diameter of the synthetic networks to match the minimum, maximum, and average values of 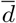 observed across the biological vasculatures (see Appendix B.1 for details on how we did so). Throughout our diameter variations, all vessel lengths and branching angles remain identical to the reference network. Therefore, Equation (12) does not apply in these cases.

Variations in 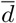 do not result in variations in 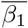 within a network. Therefore, we tracked 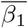 as we pruned the forking network and measured the corresponding changes in perfusion. In effect, we considered every vessel removal as a way of generating a new network with a different 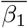.

#### 3.1.2 Network heterogeneity

To ensure heterogeneous diameters in the network, we fixed the diameter of one of the two daughter vessels (*d*_A_) at each bifurcation to be *α*-times (*α >* 1) thicker than the other (*d*_B_):

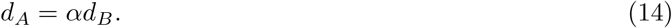

In the homogeneous case (*α* = 1.0), both daughter vessels have the same diameter. The diameter of a vessel in any generation can be calculated as:

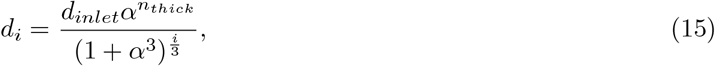

where *d*_*i*_ is the diameter of a vessel in generation *i* on a path that features *n*_*thick*_ thick vessels originating from an inlet vessel with diameter *d*_*inlet*_. Since every permutation of thick and thin vessels on a path exists in the network, it makes no difference which daughter vessel is *α*-times thicker than the other at each bifurcation. We choose the vessel that extends upwards from a bifurcation to be thicker than the lower vessel.

### 3.2 Pruning

Radiotherapy preferentially prunes smaller vessels and the extent of pruning increases with the dose [7, 8]. Therefore, we simulated increasing magnitudes of dosage by removing vessels individually in order of increasing diameter (Figure 6). Recall that the constant *λ* dictates the ratio between vessel lengths and diameters in the forking network. Therefore, pruning by diameter effectively prunes by length as well, but provides more data points than the seven distinct lengths present in the forking network.

**Figure 6:**
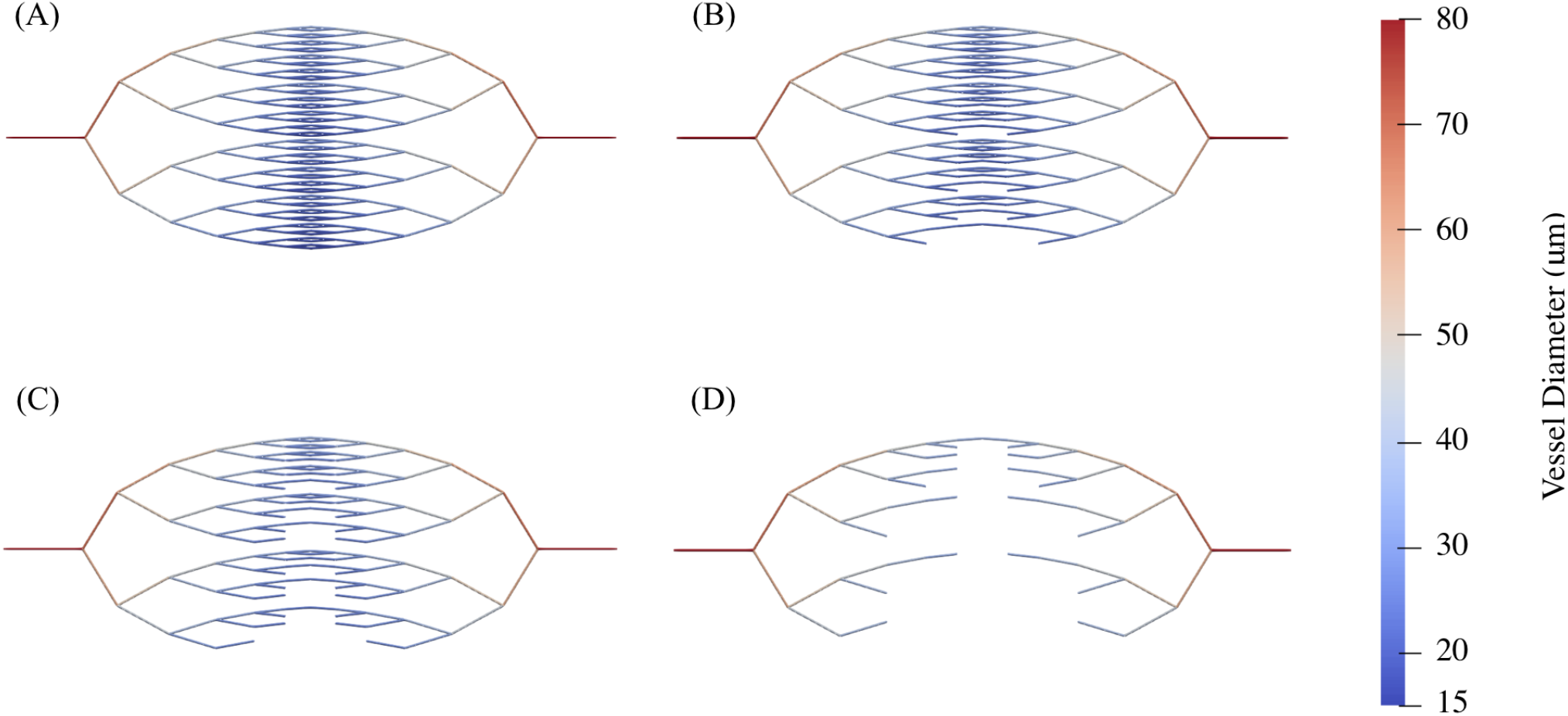
An example of pruning that shows how the forking network changes as (A) 0 vessels, (B) 50 vessels, (C) 100 vessels, and (D) 200 vessels are pruned.

Removing one vessel at a time allows us to examine pruning at a greater temporal resolution than obtainable experimentally. If two vessels have the same diameter then the vessels are pruned in order of their Vessel ID as detailed in Appendix B.2. Due to the lack of hierarchy, isolated vessels may remain in the hexagonal network during the course of pruning.

### 3.3 Perfusion threshold

Instead of identifying vessels as perfused based on fluorescence, we track the flow directly and introduce the perfusion threshold (*Q*_*min*_) as the minimum flow rate at, or above, which vessels are considered perfused and below which vessels are considered ‘hypoperfused’. This threshold serves as a proxy for the sensitivity with which our experimental apparatus can detect the signal coming from perfused vessels.

In the forking network, we set the value of this threshold *Q*_*min*_ to be 3*×* 10^*−*12^ m^3^ s^*−*1^ which, for the default model parameters and the three offsetting scenarios of the least heterogeneous network (*α* = 1.0), yields initial PFs of 0.24 (minimum), 0.50 (average), and 1.00 (maximum). In this way, we cover a large range of initial PFs spanning almost the entire interval (0, 1) which was also observed in real vasculatures (initial PFs ranging from 0.14 to 0.97). Similarly, we set *Q*_*min*_ = 3 *×*10^*−*13^ m^3^ s^*−*1^ for the hexagonal network. This value allows us to cover a sufficiently large range of initial PFs.

### 3.4 Perfusion fraction

We compute the PF using the experimental formula (Equation (1)). Guided by Equation (2), we quantify improvements in perfusion for synthetic networks by introducing Δ^max^𝒫 the maximum value attained by Δ*𝒫* during pruning, and 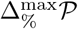 the maximum percentage change in the PF, where:

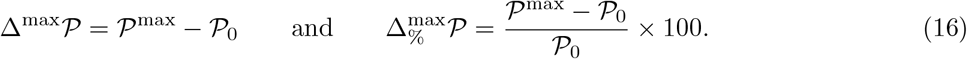

In Equation (16), *𝒫* _0_ and *𝒫* ^max^ denote, respectively, the initial and maximum value of the PF during pruning.

## 4. Results

In line with our analysis of actual tumours, we found that the largest increases in PF in our synthetic vascular networks were associated with lower values of 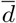 and 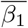. Additionally, we identified two mechanisms that can effect a positive change in perfusion post-irradiation. The vascular remodelling that occurs *in vivo* makes it difficult to isolate the effects of each mechanism. By contrast, synthetic networks offer no such barrier to analysis. We detail our inferences from these simulated networks in the following subsections.

In the absence of experimental data, we also examined the effect of diameter heterogeneity and non-hierarchical structures (modelled as hexagonal networks). We found that heterogeneity resulted in an increased and sustained Δ_%_𝒫 response, and that hierarchy was conducive to perfusion enhancement. For a full discussion, see Appendix D.

### 4.1 Two mechanisms can increase the perfusion fraction post-irradiation

One can deduce from Equation (1) that the perfusion fraction (𝒫) can increase not only if the number of perfused vessels increases after pruning, but also if the number of hypoperfused vessels decreases (Figure 7). We conclude that two mechanisms can increase the value of Δ_%_*𝒫* when a hypoperfused vessel is pruned:

**Figure 7:**
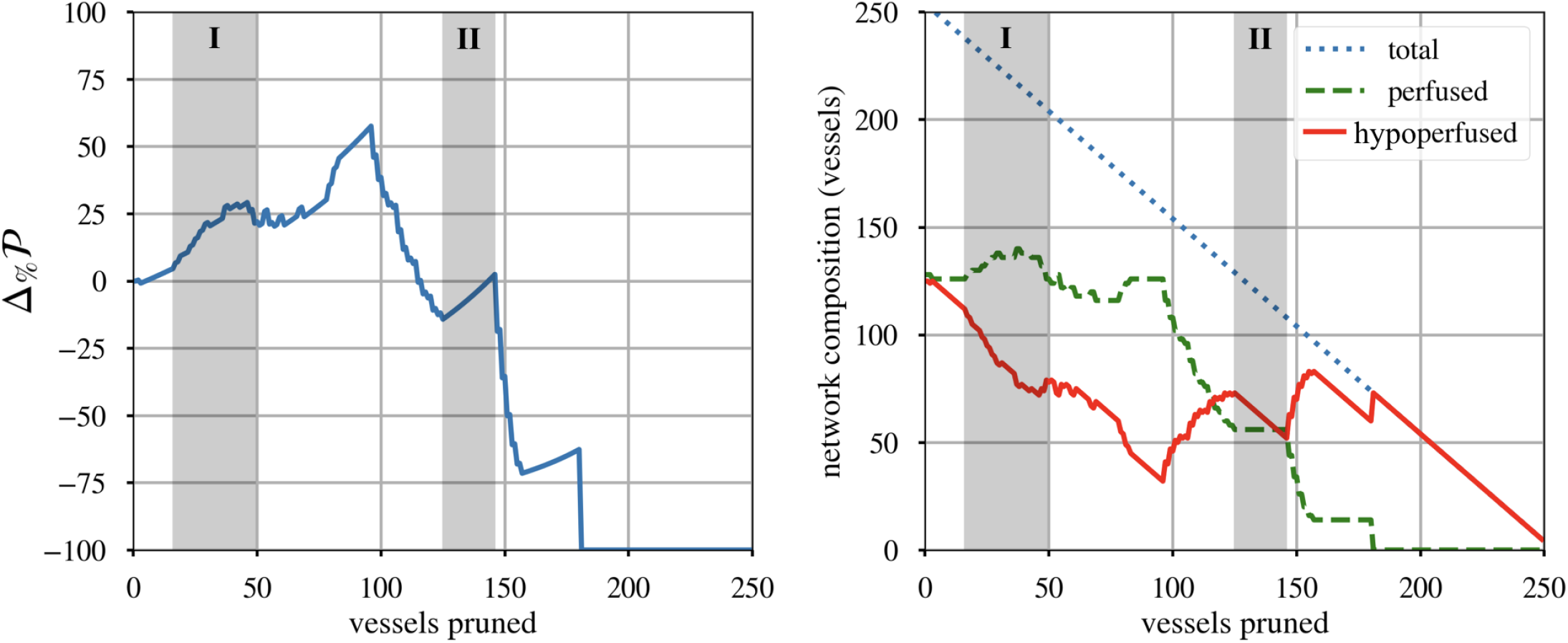
When a forking network (*α* = 1.1, 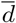 = 28.50 μm) is pruned, perfusion (Δ_%_𝒫) can increase when (I) flow is rerouted from pruned vessels to other vessels and (II) when the number of hypoperfused vessels decreases without any hypoperfused vessel becoming perfused.

- **Mechanism 1:** When a vessel is pruned, blood flow may be rerouted causing one, or more, hypoperfused vessels to become perfused and increasing the number of perfused vessels (Figure 8).
- **Mechanism 2:** Removing hypoperfused vessels increases the proportion, but not the number, of perfused vessels.

**Figure 8:**
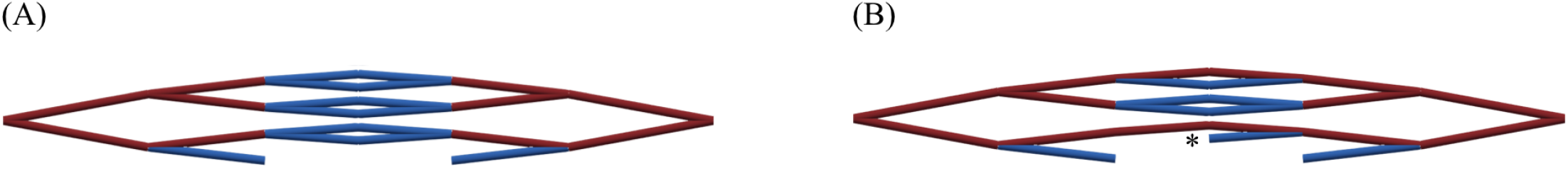
Flow is rerouted in this section of a forking network (*α* = 1.1, 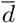 = 28.50 μm) when a single vessel is pruned (original position marked with an asterisk) between (A) and (B) and the flow rate increases in four hypoperfused vessels (blue) to the extent that they become perfused (red).

Figure 9 shows how both mechanisms act on a network during pruning. In the original, unpruned network, several hypoperfused vessels have blood flow rates that are close to the perfusion threshold while others have flow rates that are well below the perfusion threshold. When the least-perfused vessels are pruned, their flow is rerouted and the flow rate in the remaining vessels (several of which were previously hypoperfused) increases above the perfusion threshold.

**Figure 9:**
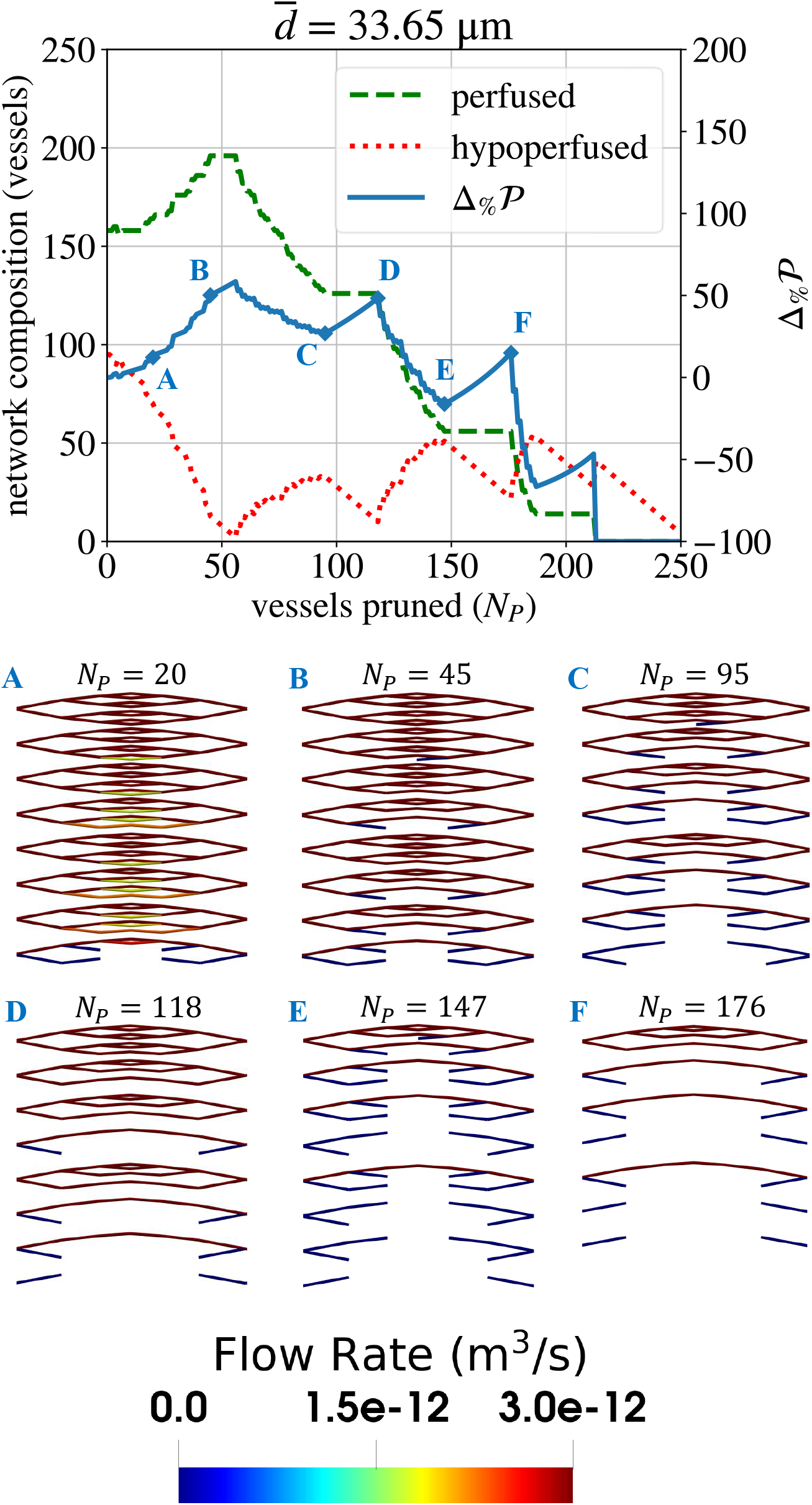
Consider the central portion of a single forking network (*α* = 1.2, 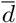). Having pruned (A) the 20 thinnest vessels, the network contains pairs of daughter vessels which experience flow rates slightly below the perfusion threshold (*Q*_*min*_ = 3 *×* 10^*−*12^m^3^ s^*−*1^). Pruning one vessel of such a pair is likely to result not only in (B) fewer hypoperfused vessels but also in more perfused vessels due to flow rerouting. Between (C) and (D) and between (E) and (F), we primarily prune blunt ends which increases the perfusion fraction (Δ_%_*𝒫*) via Mechanism 2. Between (D) and (E), Δ_%_*𝒫* dramatically decreases because perfused paths are disrupted.

A decrease in the number of hypoperfused vessels alone would correspond to a network that becomes more efficient following radiotherapy. However, it would not necessarily correspond to an improvement in oxygen distribution.

### 4.2 Mean vascular diameter determines the relative contribution of the two perfusion improvement mechanisms

Having identified two mechanisms that can increase the perfusion fraction, we next examined how the mean vascular diameter in the unpruned networks 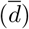 affects the percentage change in the perfusion fraction (Δ_%_*𝒫*). As in the biological experiments, we found that lower values of 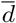 generated larger increases in Δ_%_*𝒫* and higher values of 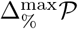 (Figure 10). However, we also examined the two mechanisms of improvement in isolation and found that higher values of 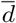 were conducive to flow rerouting, while lower values tended to improve Δ_%_*𝒫* through Mechanism 2.

**Figure 10:**
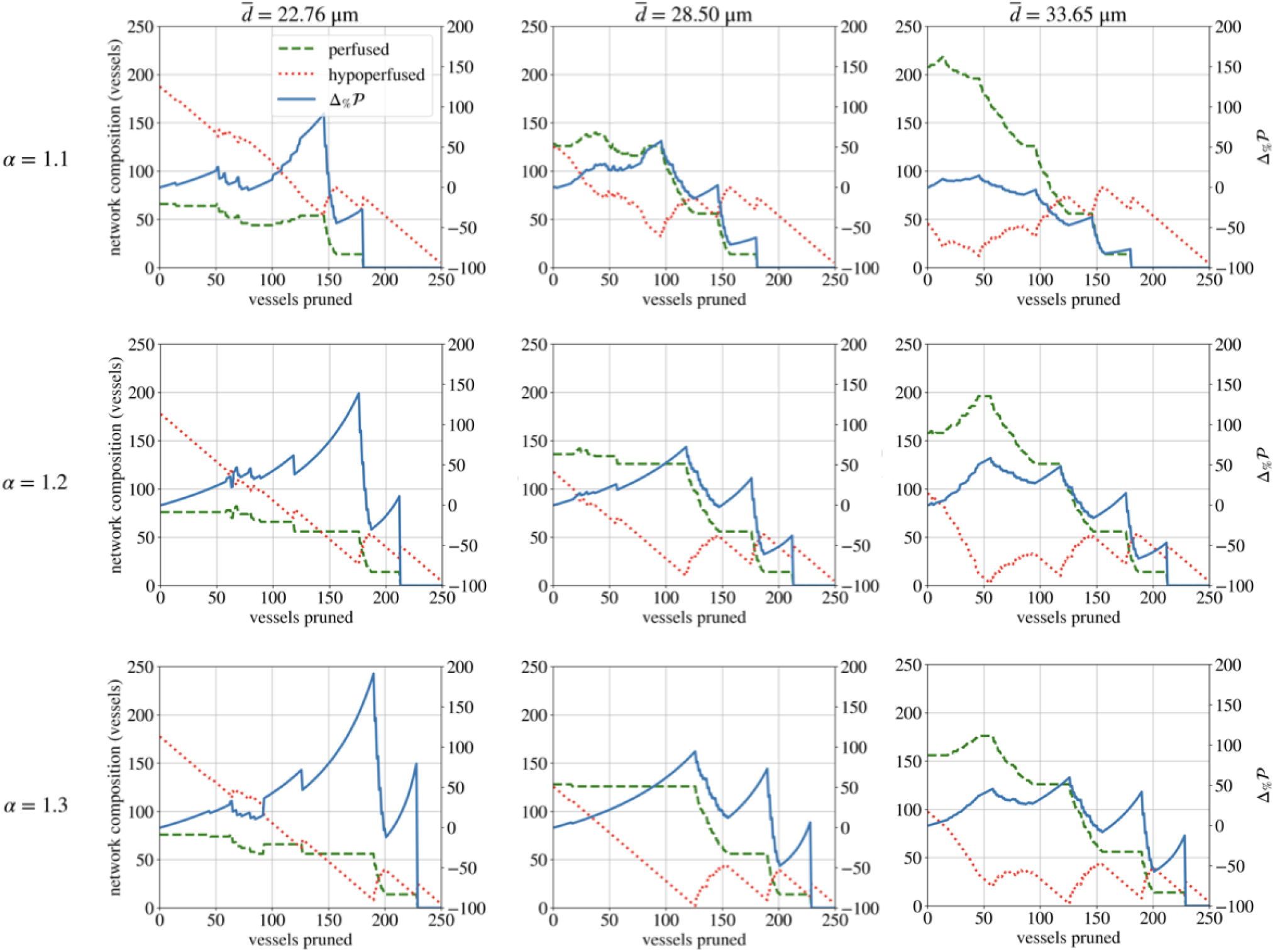
The mean diameter 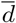 of the unpruned network influences how perfusion (Δ_%_*𝒫*) changes during pruning in the forking network across diameter heterogeneities (*α*). An increase in the number of perfused vessels is evidence of blood flow being rerouted into vessels that were previously hypoperfused. An increase in Δ_%_*𝒫* without an increase in perfused vessels is evidence of the action of Mechanism 2.

A low 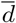 results in an increase in Δ_%_*𝒫* for two reasons. Firstly, thinner networks have lower starting perfusion fractions (*𝒫*_0_) than thicker networks (Figure 12). Therefore, any change in *𝒫* is greater relative to the network’s initial _0. 0_ is lower in thinner networks because they offer greater resistance to blood flow (Figure 11). Secondly, thinner networks have a greater proportion of hypoperfused vessels that can be pruned before any perfused vessels are pruned. In these networks, 𝒫 can increase a greater deal as a result of pruning only hypoperfused vessels.

**Figure 11:**
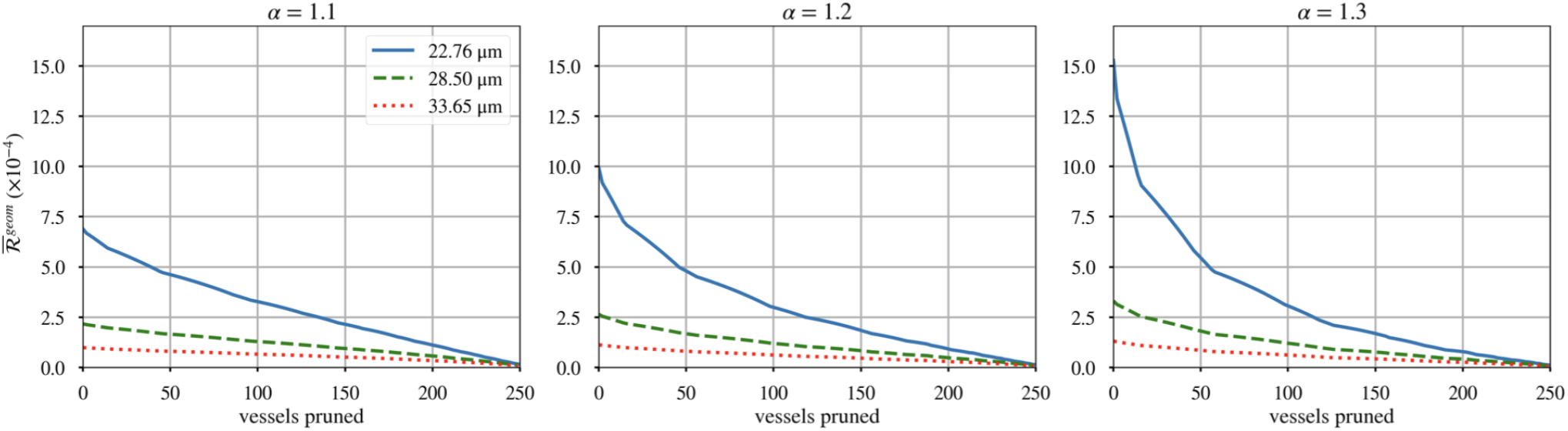
Since vessel conductivity depends on the fourth power of the mean network diameter (see Equation (3)), thinner forking networks offer substantially greater resistance 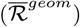 to the flow of blood. 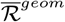 decreases monotonically during pruning because vessels are removed in order of increasing diameter. rerouting) to turn them into perfused vessels. These vessels also provide less resistance to the blood that is rerouted into them once pruning begins.

**Figure 12:**
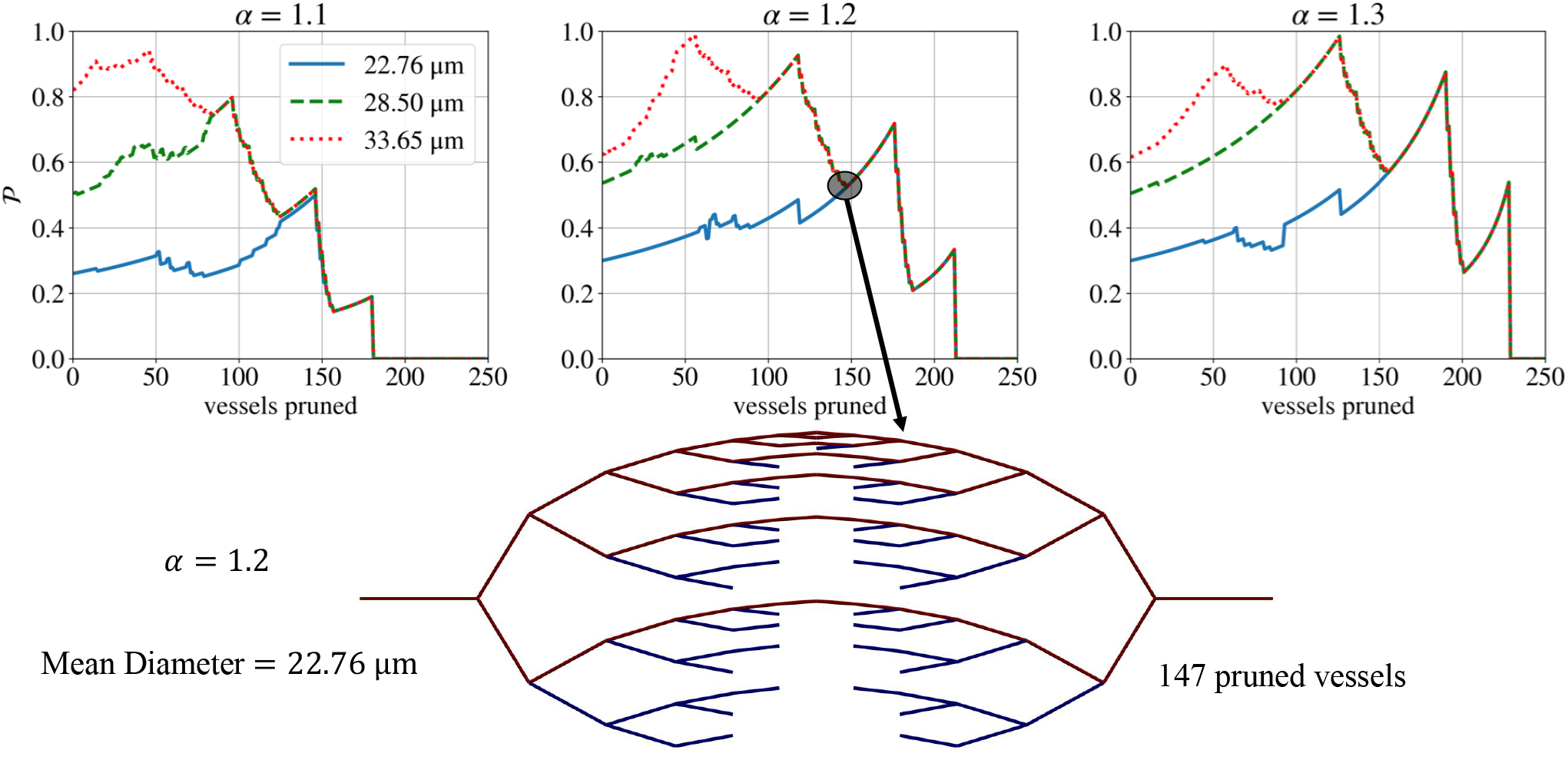
The initial perfusion fraction (*𝒫*_0_) depends on the vascular architecture (top panel). Note that for a given diameter heterogeneity (*α*), the perfusion fractions (*𝒫*) of all mean diameters 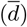 converge during the later stages of pruning, when the networks only contain thicker (low-resistance) vessels. The bottom panel shows the pruned architecture at a stage common to all values of 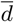 for a single value of *α*. Here, we observe that all remaining vessels are either perfused or have blunt ends (no flow). No flow rerouting can occur in the hypoperfused vessels. Thus, subsequent pruning produces the same changes in *𝒫*, regardless of the initial 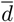. Note that the vessel colour indicates the local flow rate with the range of the colourbar adjusted so that only perfused vessels are dark red and only vessels with little to no flow are dark blue.

On the other hand, vascular architectures with a high 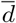 are more susceptible to increases in Δ_%_ 𝒫 through blood flow rerouting (Figure 10). For these architectures, flow rates in the pruned (hypoperfused) vessels are closer to the perfusion threshold. Therefore, these vessels require a smaller increase in blood supply (via

### 4.3 A drop in the proportion of loops precedes a rise in perfusion

Thus far, we have compared networks that differ in terms of their initial, unpruned architecture. Since pruning produces blunt ends (thereby changing the number of loops as discussed in Section 2.2.3), the pruned networks can serve as proxies for networks that initially contain different proportions of angiogenic sprouts. Contrasting the number of loops per vessel 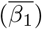 with the change in perfusion fraction (Δ_%_𝒫), we observed two distinct results (Figure 13).

**Figure 13:**
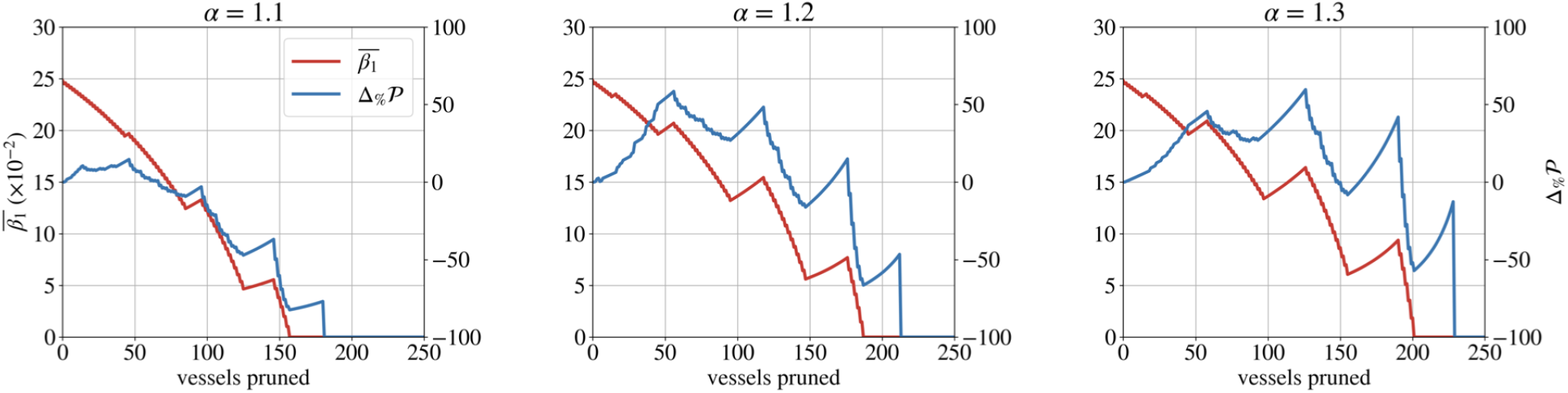
A drop in the number of loops per vessel 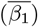 precedes a positive change in the perfusion fraction (Δ_%_𝒫) in the forking networks 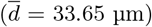 Minor differences in 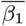 arise between heterogeneities because *α* changes the order of pruning by virtue of changing the diameters of vessels.

In the early stage of pruning (less radiation-induced cell death), Δ_%_*𝒫* typically increases regardless of the behaviour of 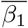. In this stage, pruning removes the thinnest vessels, which tend to constitute hypoperfused loops or blunt ends. Thus, Δ_%_*P* increases through both mechanisms of improvement.

In the later stages of pruning (more radiation-induced cell death), however, peaks and troughs in 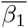 correspond to those in Δ_%_ 𝒫 In particular, a reduction in 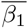 often precedes an increase in Δ_%_𝒫. When a vessel that separates two loops is pruned, the loops merge into one and the total number of loops decreases. At the site of the pruned vessel, a blunt end may be created. Subsequent pruning favours the removal of this blunt end due to its small diameter by virtue of previously being connected to the smallest vessel (recall Murray’s Law, Equation 11). Pruning the blunt-ended vessel does not change the total number of loops. However, it increases the proportion of loops 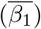 and the resulting reduction in the hypoperfused vessel count leads to an increase in Δ_%_𝒫. Therefore, a decrease in 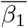 indicates that the network now contains a greater proportion of blunt ends (and vice-versa).

Thus, we find that Δ_%_*𝒫* increases when networks with a low value of 𝒫are pruned. We also infer that 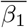 indicates when an improvement seen in Δ_%_𝒫 is caused by the pruning of blunt-ended vessels with no flow, rather than by rerouting.

## 5. Discussion

In this paper, we used a computational model to investigate the influence of vascular architecture on the outcome of radiation-induced pruning. As per [8], we measured these outcomes in terms of overall network perfusion, as defined by the PF (Equation (1)).

In our experimental data, we found that perfusion typically improved in vascular networks with lower mean diameters 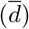 and fewer loops per vessel 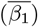. It may seem counter-intuitive for network perfusion to increase after vessels are removed in a model that excludes angiogenesis. However, our synthetic networks mirrored our observations of the experimental data. Using our *in silico* model, we elucidated the mechanisms underlying these correlations, including two mechanisms of improvement in the PF.

We have also shown that different architectural features are susceptible to different mechanisms of improvement in the PF. In particular, we found that networks with low average diameters tend to exhibit improvements in PF via the reduction of hypoperfused vessels, while networks with large average diameters are more prone to rerouting (Section 4.2). In the former, large doses of radiotherapy (i.e. more vessels pruned) seem to improve the PF, while rerouting seems to occur in the latter with smaller doses of radio-therapy (Figure 10).

Therefore, our model may explain the differential perfusion responses reported in the literature [9]. Moreover, our exposition of the different mechanisms of PF improvement may explain why Bussink et al [12] observed an increase followed by a decrease in tumour perfusion. Bussink et al. attributed the increase in perfusion to the redistribution of blood and the refilling of previously non-functional vessels (in line with our model’s predictions).

### 5.1 Experimental implications

Our findings also have several implications for experimental methodologies used to study radiation-induced pruning. For one, our results in Section 4.1 have shown that the PF is not an ideal measure of a tumour’s perfusion, since it fails to account for hypoperfused vessels that are pruned. Furthermore, the PF offers no indication of a vascular network’s ability to sufficiently deliver oxygen in tissue. Theoretically, a region could feature a single perfused vessel and have a PF of 1 despite the fact that the vessel may not cover a sufficient area or carry enough red blood cells to maintain a normoxic environment. Experimentally, Bussink et al. found discrepancies between perfusion and hypoxia [12].

Monitoring the PF alone cannot tell us which mechanism (as discussed in Section 4.1) is at work during pruning. Discerning the mechanism is important because it tells us whether or not blood flow has improved in the remaining vessels. In Section 4.3, we showed that the number of loops per vessel may act as an indicator of improvements in PF via the reduction of hypoperfused vessels.

In Table 2, we classified biological tumours into groups A and B based on the Δ 𝒫on a fixed day. However, the non-monotonic response of 𝒫in our simulations implies that this classification is subject to the sampling point (Table 4). In other words, a network belongs to group A or B depending on the number of vessels pruned from it.

**Table 4:** The classification of forking networks into groups A and B varies based on the sampling point, i.e. how many vessels have been pruned when Δ_%_*𝒫* is measured. Cells have been shaded green or red to represent positive or negative values of Δ_%_*𝒫*.

| Dosage (vessels killed) | $\bar{d} = 22.76 \mu\text{m}$ | | | $\bar{d} = 28.50 \mu\text{m}$ | | | $\bar{d} = 33.65 \mu\text{m}$ | | |
| --- | --- | --- | --- | --- | --- | --- | --- | --- | --- |
| | $\alpha = 1.1$ | $\alpha = 1.2$ | $\alpha = 1.3$ | $\alpha = 1.1$ | $\alpha = 1.2$ | $\alpha = 1.3$ | $\alpha = 1.1$ | $\alpha = 1.2$ | $\alpha = 1.3$ |
| 25 | 7.13 | 10.48 | 10.48 | 15.66 | 12.11 | 8.75 | 8.36 | 16.07 | 13.31 |
| 50 | 20.26 | 24.02 | 20.76 | 22.08 | 22.20 | 22.08 | 8.52 | 53.85 | 39.92 |
| 75 | -1.49 | 37.62 | 15.30 | 28.09 | 30.95 | 39.13 | -6.23 | 39.55 | 32.28 |
| 100 | 9.52 | 42.67 | 42.67 | 38.62 | 52.21 | 61.72 | -14.70 | 31.01 | 32.69 |
| 125 | 60.46 | 44.51 | 70.32 | -14.20 | 41.32 | 93.06 | -47.20 | 21.65 | 58.41 |
| 150 | 17.95 | 79.25 | 79.25 | -35.38 | 0.17 | 21.63 | -60.23 | -13.78 | -0.20 |
| 175 | -32.07 | 135.98 | 135.98 | -64.97 | 31.87 | 40.11 | -78.44 | 13.51 | 14.96 |
| 200 | -100.00 | -13.69 | -1.36 | -100.00 | -51.77 | -41.44 | -100.00 | -58.49 | -51.95 |
| 225 | -100.00 | -100.00 | 60.71 | -100.00 | -100.00 | -4.58 | -100.00 | -100.00 | -21.71 |

Finally, we also computed for forking networks a compound metric discussed in Section 2.2.4: the resistance per loop. As expected from Equation (10), over the course of pruning, the trend of 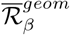follows that of 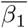 in an inverse fashion (Figures 13 and 14). However, 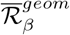 also encodes information on the mean diameter (via resistance) and thus allows us to distinguish between networks of varying mean diameters. Consistent with real vasculatures, we observe that networks with initially higher 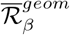 exhibit greater potential for perfusion improvement, as measured by 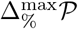 (compare with Figure 10). This leads us to propose the metric 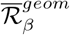 for future investigations as a predictor of perfusion improvement, useful especially when 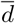 and 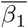 individually give contradictory predictions. This might be relevant, for example, for vasculatures with low mean diameters and a large proportion of loops, even though such a combination of characteristics did not occur within the small tumour sample studied here. These speculations need to be confirmed by future work and the mechanism of improvement must also be discerned.

**Figure 14:**
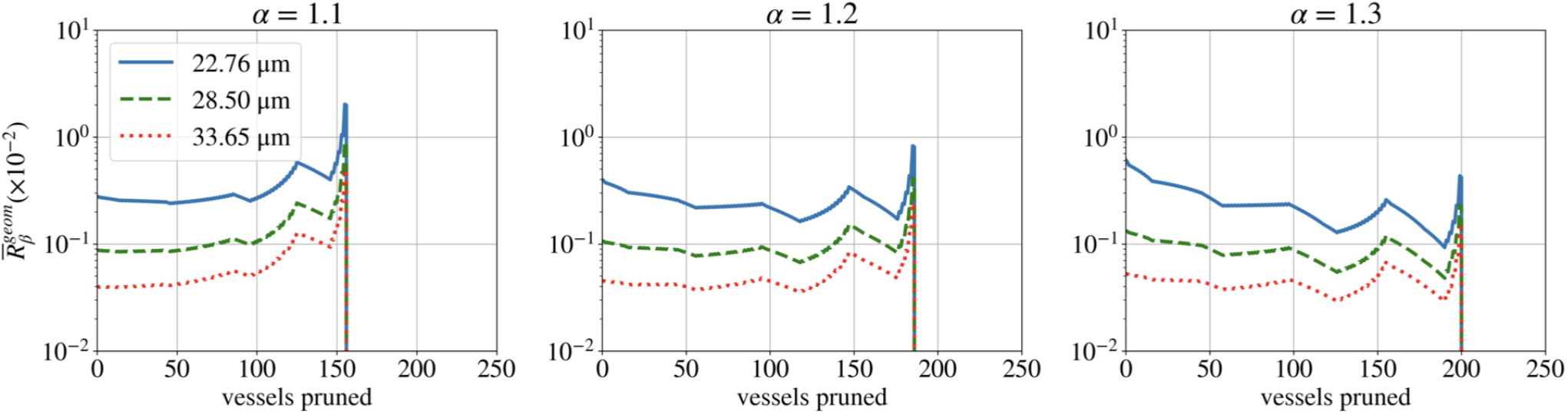
The resistance per loop follows trends inverse to those observed for loops per vessel (Figure 13), but can also distinguish between networks of varying mean diameter.

### 5.2 Future work

In this work, we used forking and hexagonal networks to illustrate the role of hierarchy in vascular architecture. However, both network types are regular (i.e. symmetric) and might not adequately represent all features of dysregulated tumour vasculature. As a next step, one should consider irregular networks, such as those generated by the Voronoi tessellation of a planar region [46, 47].

We performed our blood flow simulations under the assumption that the haematocrit (fraction of blood made up of red blood cells) distributes itself evenly in all vessels. In reality, however, the spread of haematocrit at vessel junctions is more complex. The haematocrit in each vessel affects its flow rate (Equation 17), as well as its capacity to carry oxygen. Additionally, our colleagues have hypothesised that irradiating a tumour with a larger proportion of small vessels might be more likely to lead to improved perfusion by altering the proportion of haematocrit splitting [8]. Therefore, it will be important to contrast different haematocrit splitting rules sourced from the literature [13, 48, 49].

Different splitting rules will give rise to different oxygen distributions in the tissue. These distributions are a key determinant of the efficacy of standard oncological treatments, such as chemotherapy, which can also be modelled within Microvessel Chaste [43, 44] as a chemical transport process. The death of tumour cells caused by radiotherapy or chemotherapy lowers net oxygen consumption, which further increases tissue oxygen levels. The inclusion of an oxygen consumption term proportional to the local density of live tissue cells will be an important next step in the precise modelling of tissue oxygen perfusion.

We have already highlighted that the PF is an imprecise metric to describe the oxygenation status of the tissue. Once we explicitly simulate oxygen diffusion, we will need to adopt and develop new spatial metrics that better describe the spread of oxygen.

With a model of oxygenation in place, we will then consider the dynamic nature of the tissue perfused by the vasculature. Cells that comprise the tissue will have varying uptake rates, and will divide and die based on the amount of oxygen available to them.

Let us note that the existing model can also be adapted to better represent empirical radiation killing by pruning vessels in order of increasing flow rate, thus modelling the regression of hypoperfused vessels as observed in developmental vascular networks [31, 32]. Pruning can also be conducted stochastically and modified to reflect radioresistance when the region is anoxic.

Since the vasculature itself is also dynamic, future models must factor in structural adaptation and the secretion of angiogenic factors. Random sprouting models may be useful here [50]. Percolation models would also serve as a good representation of the statistical distribution of avascular spaces found in tumours [51]. Moreover, percolation models have been used to replicate observed oxygenation conditions in mice tumour xenografts [51]. Such a study may also involve modifying haematocrit splitting rules to include higher-order splitting, since tumour vasculature is not limited to bifurcations [13].

Finally, we may leverage our insights from synthetic networks and apply them to a real biological network obtained via the methods outlined in [22].

## 6 Materials and methods

### 6.1 Experimental procedures and data preprocessing

#### 6.1.1 Experimental procedures

##### Abdominal imaging window implantation

This procedure was based on a previously described method [52]. Transgenic mice with fluorescent protein tdTomato expressed in endothelial cells on C57Bl/6 background were prepared in a surgical unit, and administered with inhalational anaesthesia and pre-operative analgesics. Body temperature and respiration rates were monitored throughout the procedure. A 1 cm cut was made along the abdominal midline approximately 5 mm underneath the sternum followed by blunt dissection around the cut to separate the connective tissue from the skin. A custom-made imaging window frame (Workshop at the Department of Oncology, Oxford University) was fitted underneath the skin. Continuous sutures were used to secure the skin to the window frame. Approximately 2.5 *×*10^5^ MC38 cells (murine colon adenocarcinoma cells) stably expressing eGFP in 5 μL containing 30% of Matrigel and 10% of Evan’s blue dye were injected under the connective tissue and above the abdominal muscle layer. The chamber was then flushed with water to lyse non-injected cells by osmotic shock, tapped dry with sterile cotton swabs and flooded with saline. A cover glass glued on the chamber’s lid was secured onto the window frame. The animals were then placed onto a heat mat for post-operative recovery, and their health and tumour growth was monitored by visual examination. All animal experiments were conducted in accordance with the United Kingdom Animals (Scientific Procedures) Act 1986 as amended (Amendment Regulations 2012 [SI 2012/3039]), under the authority of a UK Home Office Project License (PPL 30/2922 and PCDCAFDE0), with local ethical approval from the University of Oxford Animal Welfare and Ethical Review Panel.

##### Treatment regimes

Animals with tumours approximately 100 mm^3^ growing in the window chamber were administered radiation treatment. Prior to the radiation treatment, mice were anaesthetised under inhalation with isoflurane and placed in an imaging-guided small animal radiation research platform (SARRP) irradiator (Xstrahl Ltd). A Cone Beam CT scan (computerised tomography) of each mouse was obtained and the treatment was planned using Muriplan (Xstrahl Ltd). The SARRP was used to deliver 15 Gy of X-rays (220 kVp copper filtered beam with HVL of 0.93 mmCu) to the tumour at 2 Gy per minute. Dosimetry of the irradiator was performed as previously described [53]. We refer to the start of treatment as Day 0.

##### Intravital two-photon imaging

Mice were imaged for seven days after radiation treatment with a Zeiss LSM 880 microscope equipped with a respiratory monitoring system. The stage and atmosphere were heated to 37 ^*?*^C. To label perfused vessels, Qtracker™ 705 Vascular Labels (0.2 μM, ThermoFisher Scientific) were infused intravenously using a motorised pump at a rate of 0.84 μL·min^*−*1^. A mode-locked MaiTai laser tuned to 920 nm was used to simultaneously excite eGFP, tdTomato, and Qtracker705. The Qtracker705 signal was acquired through a BP700/100 filter with a non-descanned detector. GaAsP detectors were used to acquire the signal of tdTomato selected by a BP 650/45 filter and the eGFP selected by a BP525/50 filter. Images were acquired in Z-stack tile scans with a pixel size of 0.823 μm and an image size per tile of 512*×* 512*×* 5 in *x, y*, and *z*, respectively. A water immersion 20*×* objective made for UV-VIS-IR transmission with a numerical aperture of 1.0 was used. Representative examples of vascular networks from our experiments are displayed in Figure 2.

#### 6.1.2 Data preprocessing

The biological networks were obtained by multiphoton intravital 3D imaging [54] and consisted of 3D stacks of images of tumour blood vessels. Skeleton files were extracted from the imaging data by combining two segmentation models and taking their geometric average. The skeletons were then pruned (see reference [55], p. 165, for a full description). We extracted blood vessel networks from skeleton files using the method VesselTree from unet core.vessel analysis in the Python code package unet-core [56]. The extracted networks consist of points on vessel branches (multiple points per vessel branch including branching points) which represent the network nodes, and the vessels that connect them which constitute the edges of the network. VesselTree also enables us to extract network features such as vessel diameters and lengths.

### 6.2 Simulation methods

#### 6.2.1 Blood flow

We assume that the blood flow rate *Q* in a vessel of length *L* and diameter *d* is determined by Poiseuille’s law (Equation (3)). Following [13, 36], the effective blood viscosity in Equation (3) is expressed as:

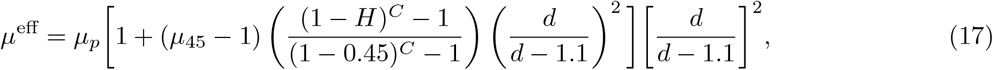

where *H* is the discharge haematocrit, *d* is the vessel diameter, *μ*_*p*_ is the plasma viscosity,

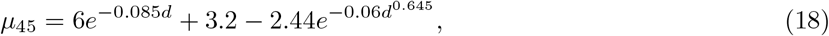

and

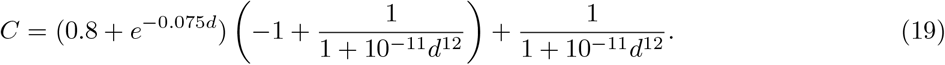

As discussed in Section 2.2.2, we assume that haematocrit is distributed evenly within the network (*H* = *H*_*inlet*_ = 0.45) to yield a linear flow problem. With signed flow rates *Q*_*i*_ for each vessel *i* (i.e. each edge connected to a node), we also impose conservation of blood at each network bifurcation node:

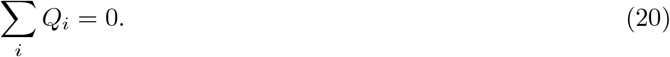

In each (unpruned) network, the leftmost nodes serve as inlets and all other vessels with one detached node serve as outlets. Inlet nodes are assigned a pressure (*p*_*inlet*_) of 3333 Pa (*≈* 25 mmHg), while outlet nodes are assigned a pressure (*p*_*outlet*_) of 2000 Pa (*≈* 15 mmHg) [57].

#### 6.2.2 Software development and post-processing

We performed our simulations using Microvessel Chaste (version 3.4.3), an add-on to the open-source simulation package Chaste (version 2020.1) [43, 44]. We added custom functionality to the base version of Microvessel Chaste, including new network generators and pruning functions. We generated synthetic networks detailed above and pruned them vessel-by-vessel in order of increasing vessel diameter. At each pruning step, we updated the flow distribution, and exported the perfusion fraction and the network itself in the .vtk format, accounting for all vessels that had not yet been pruned.

We used Python scripts to generate from the .vtk files adjacency matrices weighted by vessel diameter and vessel length, from which we calculated key geometric metrics, such as mean diameter and mean geometric resistance. Based on a MATLAB package [58], we wrote Python scripts to import the adjacency matrices from above, calculate both the number of loops and loops per vessel at each pruning stage using Equation (6), and finally divide the mean resistance by the latter.

## Data and Code Availability

### Experimental data

Experimental data used in this study have been published in [8] and can be found at [33]. Post-processing files relating to these vasculatures are available from the Zenodo database (DOI https://doi.org/10.5281/zenodo.8011700). All other experimental data needed to evaluate the conclusions in the paper are within the manuscript.

### Synthetic networks simulations

The modified Microvessel Chaste source code, along with the Python files used in post-processing, can be found on GitHub (github.com/vedangnarain/enhanced-perfusion-following-exposure-to-radiotherapy.git).

### Topological data analysis

Post-processing files calculating topological metrics considered in this manuscript can be found on GitHub (github.com/vedangnarain/enhanced-perfusion-following-exposure-to-radiotherapy.git).

## A Reliability of the data as measured by the number and size of connected components

As noted in Section 2.2, the number of vessels for tumour vasculature 7 increased throughout the first four days post-irradiation. Table 5 documents that this vasculature had, relative to the network size, an extremely large number of connected components (CC) and an extremely small largest connected component, indicating problems in image processing, which is why this tumour was excluded from subsequent analysis.

**Table 5:** Statistics relating to the number and size of connected components indicate problems with image acquisition for tumour number 7.

| **Tumour statistics on Day 0** | **1** | **2** | **3** | **4** | **5** | **6** | **7** |
| --- | --- | --- | --- | --- | --- | --- | --- |
| Vessel count (network size) | 1000 | 4087 | 1038 | 2175 | 644 | 528 | 436 |
| Number of CCs $\beta_0$ | 301 | 184 | 173 | 301 | 82 | 86 | 184 |
| Number of CCs per size $\bar{\beta}_0$ | 0.30 | 0.05 | 0.17 | 0.14 | 0.13 | 0.16 | 0.42 |
| Size of largest CC | 168 | 3467 | 317 | 1365 | 338 | 353 | 59 |
| Size of largest CC per size | 0.17 | 0.85 | 0.31 | 0.63 | 0.52 | 0.67 | 0.14 |

## B Fitting the forking network and assigning vessel IDs

### B.1 Adjusting vessel diameters to match summary statistics for real vasculatures

We fix the number of generations at 7 throughout this work so that the vessel diameters span a sufficiently large range. In order to generate significant intra-generational diameter heterogeneity we consider three values of *α*, namely 1.1, 1.2, and 1.3. Considering the smallest and the largest *α*, the means and standard deviations of the diameter distributions lie within the ranges delineated by summary statistics for the real vasculatures if and only if the inlet diameter is roughly in the range 80 *−*100 μm (see Figure 15).

**Figure 15:**
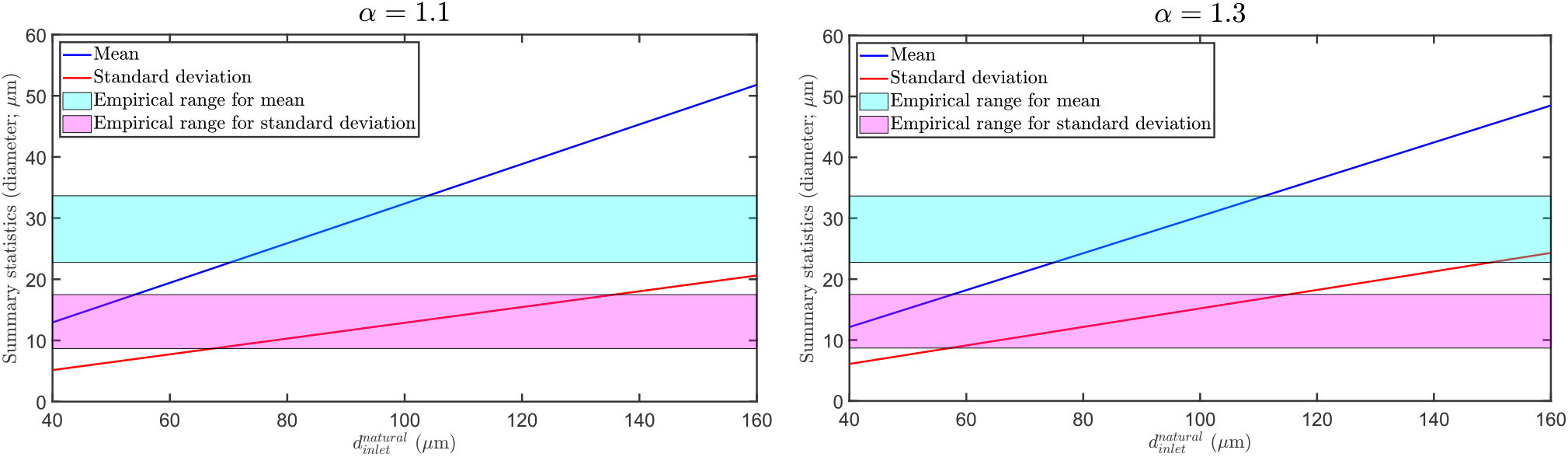
The means (blue) and standard deviations (red) of diameter distributions in Murray’s law forking networks with 7 generations of vessels as a function of the inlet diameter 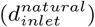 for *α* = 1.1 (left panel) and 1.3 (right panel). The coloured areas delineate ranges of means (cyan) and standard deviations (magenta) of diameter distributions found in real vasculature.

Taking the inlet diameter to be 90 μm, the top panels in Figure 16 show bar charts documenting the diameter distributions for varying *α*, while the bottom panels detail how the diameters are distributed within individual generations.

**Figure 16:**
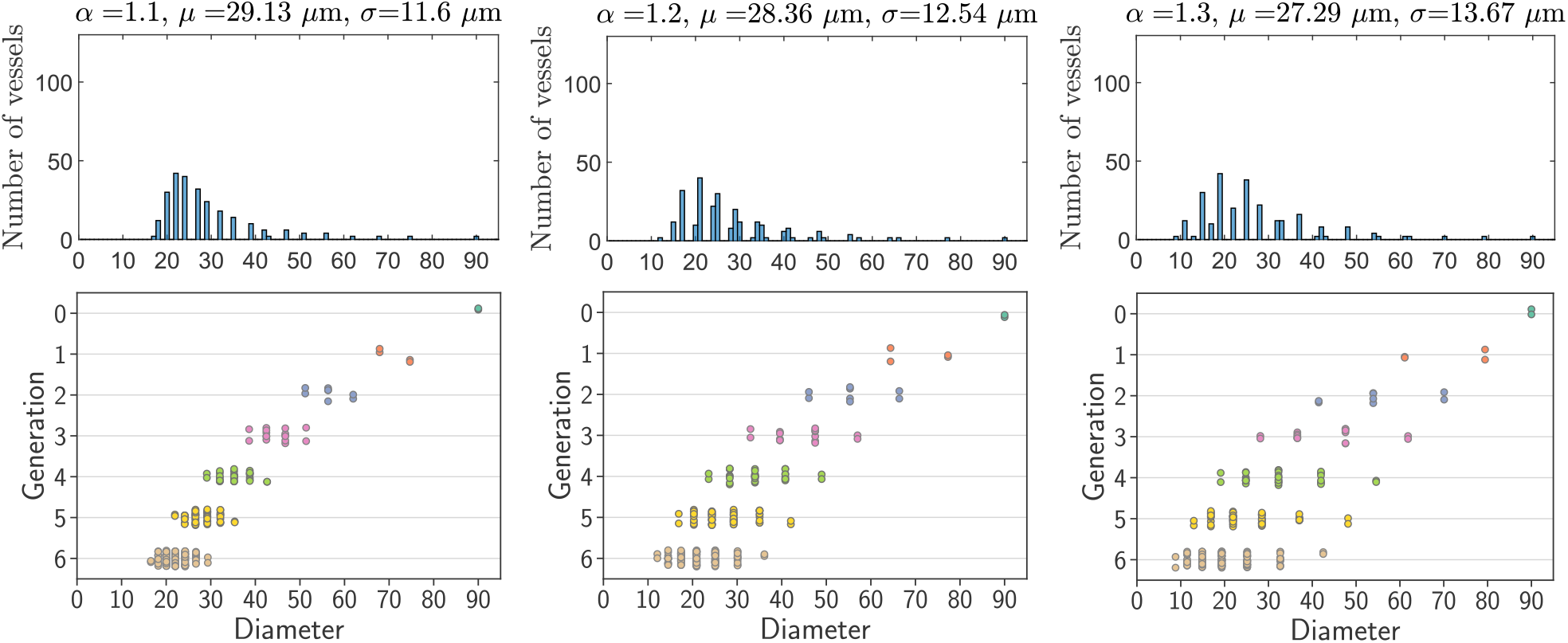
The parameter *α* dictates the diameter distribution of the Murray’s law forking networks.

Note from Figures 15 and 16 that both increasing the inlet diameter for a fixed *α* and increasing *α* for a fixed inlet diameter lead to an increase in standard deviation. However, both approaches also yield an altered mean diameter. As we wish to study the impact of the mean and the standard deviation on network perfusion (as affected by radiotherapy-induced vessel pruning) independently, we proceed in a different manner.

We aim to find a new value for the inlet diameter to obtain realistic standard deviations for all studied values of *α*. We also aim to offset the diameters of all vessels in the Murray’s law forking networks (i.e. networks obeying Equation (11)) by the same constant to match the mean diameter in real vasculatures, namely the smallest (22.75 μm), average (28.5 μm), and largest (33.65 μm) mean diameters. Following Figure 16, this simply means that all data points in strip plots and all columns in histograms will be shifted right or left to match the means observed in real data. Note that this offsetting does not alter the standard deviation. On the other hand, for a fixed offset, *α* only modulates the standard deviation.

After specifying the number of generations, *α* and the offset, all vessel diameters follow from the inlet diameter. To limit the possible range for the inlet diameter, we require that for all three values of *α*, the Murray’s law forking networks manifest standard deviations within the experimentally-observed range (from 8.68 μm for the least to 17.49 μm for the most heterogeneous vasculature). From Figure 15, we thus conclude an approximately 70*−*110 μm range for the inlet diameter of the Murray’s law forking network.

Finally, while we have designed our offset forking networks so that they match the means and standard deviations found in real vasculatures, we must make sure that we do not assign any unrealistically small — or even negative — diameters (for all values of *α*). In Figure 17, we plot the minimum vessel diameter as a function of the inlet diameters for two extreme values of *α* (1.1 and 1.3) and for the three offsets used in our simulations. The value *d*_*inlet*_ = 100 μm used in [13] would result (using *α* = 1.3 and offsetting to match the smallest mean diameter in real vasculatures) in vessel diameters well below both the smallest diameter found in our biological networks (*≈*7.39 μm) and even the minimum diameter of an undeformed red blood cell, i.e. 6 μm [59]. Therefore, we will use *d*_*inlet*_ = 75 μm as default in the Murray’s law forking networks, which yields plausible vessel diameters in all scenarios.

**Figure 17:**
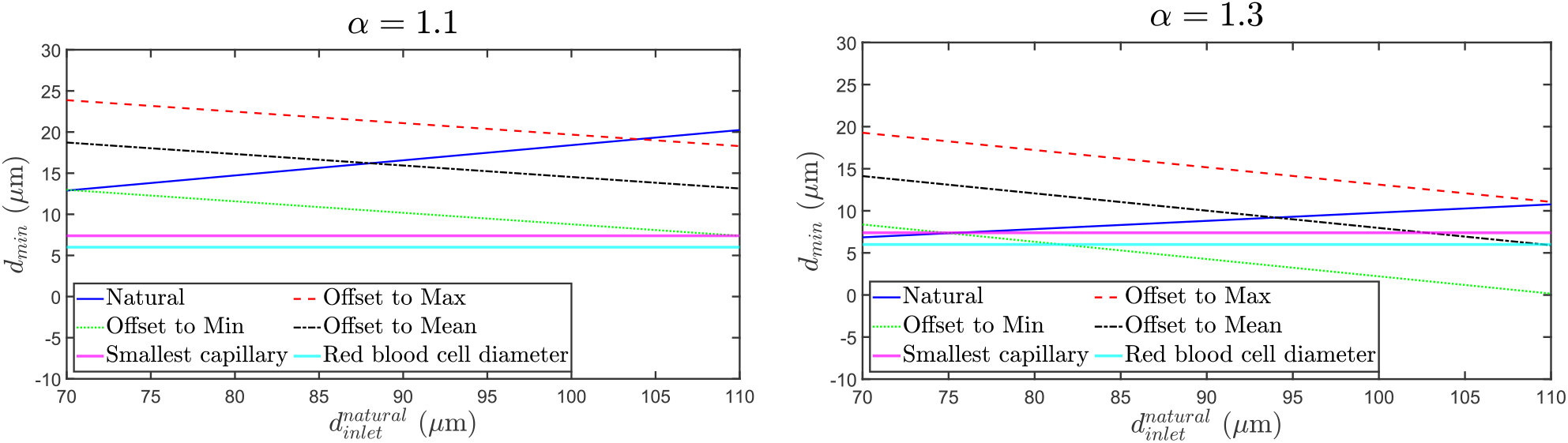
The blue curves show minimum vessel diameter (*d*_*min*_) as a function of inlet diameter 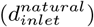 in Murray’s law forking network with seven generations of vessels for *α* = 1.1 (left panel) and 1.3 (right panel). The minimum diameter changes due to offsetting to match the smallest (dotted green), average (dash-dotted black), and largest (dashed red) mean diameter in real vasculatures. The diameter of the smallest vessel found in real vasculatures (*≈* 7.39 μm) is indicated with magenta and the minimum diameter of red blood cells (6 μm) with cyan lines.

### B.2 Assigning vessel IDs

During vessel network generation in Microvessel Chaste, we assign each vessel a unique ID which is an integer number that will be used to determine the order of pruning in case two or more vessels have the same diameter. The way in which vessel IDs are assigned is illustrated in Figure 18. In the left (diverging) half of the domain, vessels are assigned IDs that increase with increasing generation number and their mirror images in the right (converging) half of the domain are always assigned IDs equal to those of their left-half counterpart plus 2.

**Figure 18:**
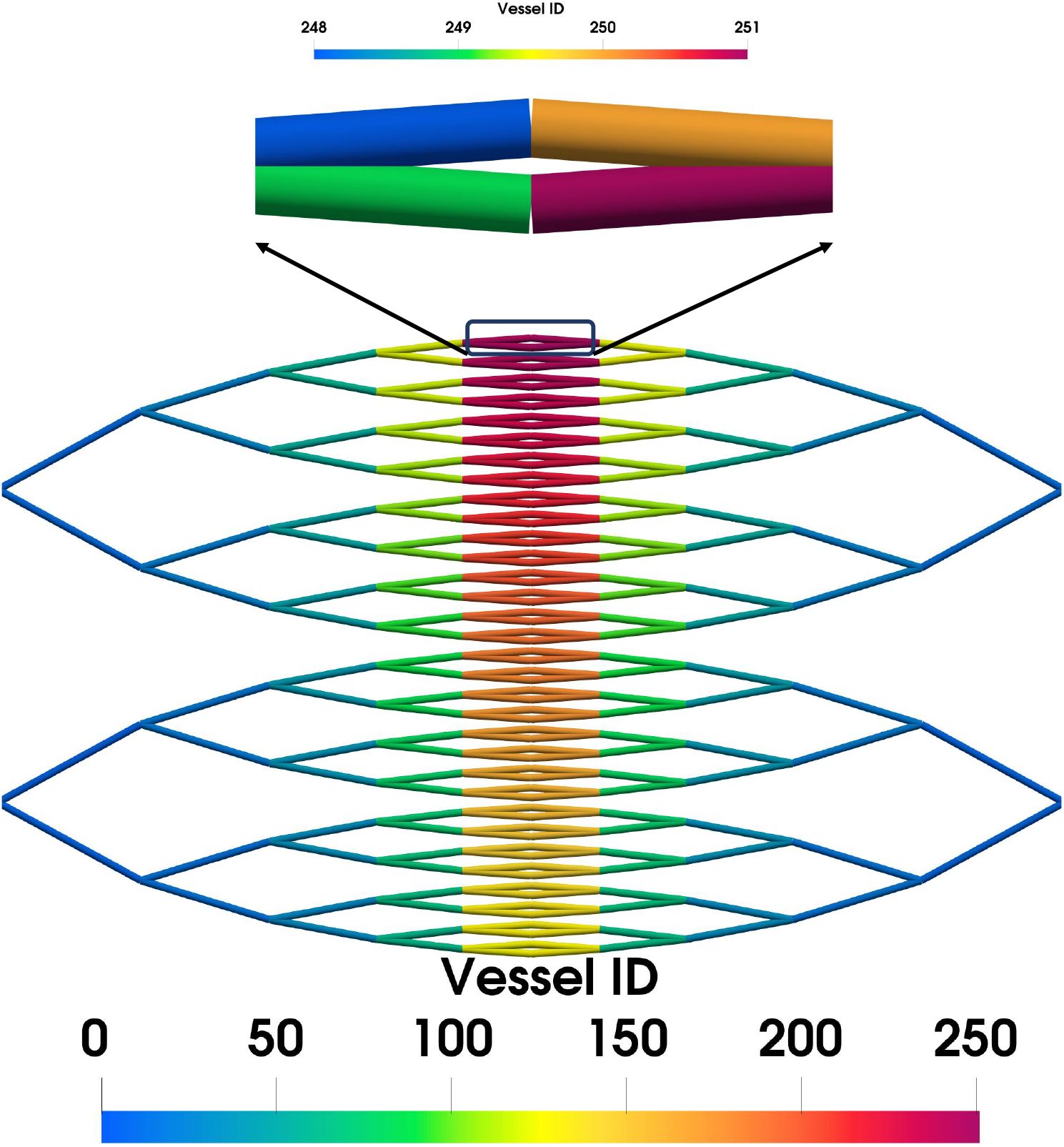
Vessel IDs are assigned based on the order in which the vessels are added to the network during network generation, as demonstrated here. Note that the range of the colourbar in the top panel has been adjusted to demonstrate the vessel IDs assigned for two daughter vessels of the same parent vessel as well as for their mirror images in the right (converging) part of the domain.

## C Design of hexagonal network

In the non-hierarchical network, regular hexagons populate a two-dimensional domain measuring 2 mm by 2 mm (Figure 19). Each vessel has a length (*L*_*hex*_) of 100 μm. The network has 11 inlets and 10 outlets.

**Figure 19:**
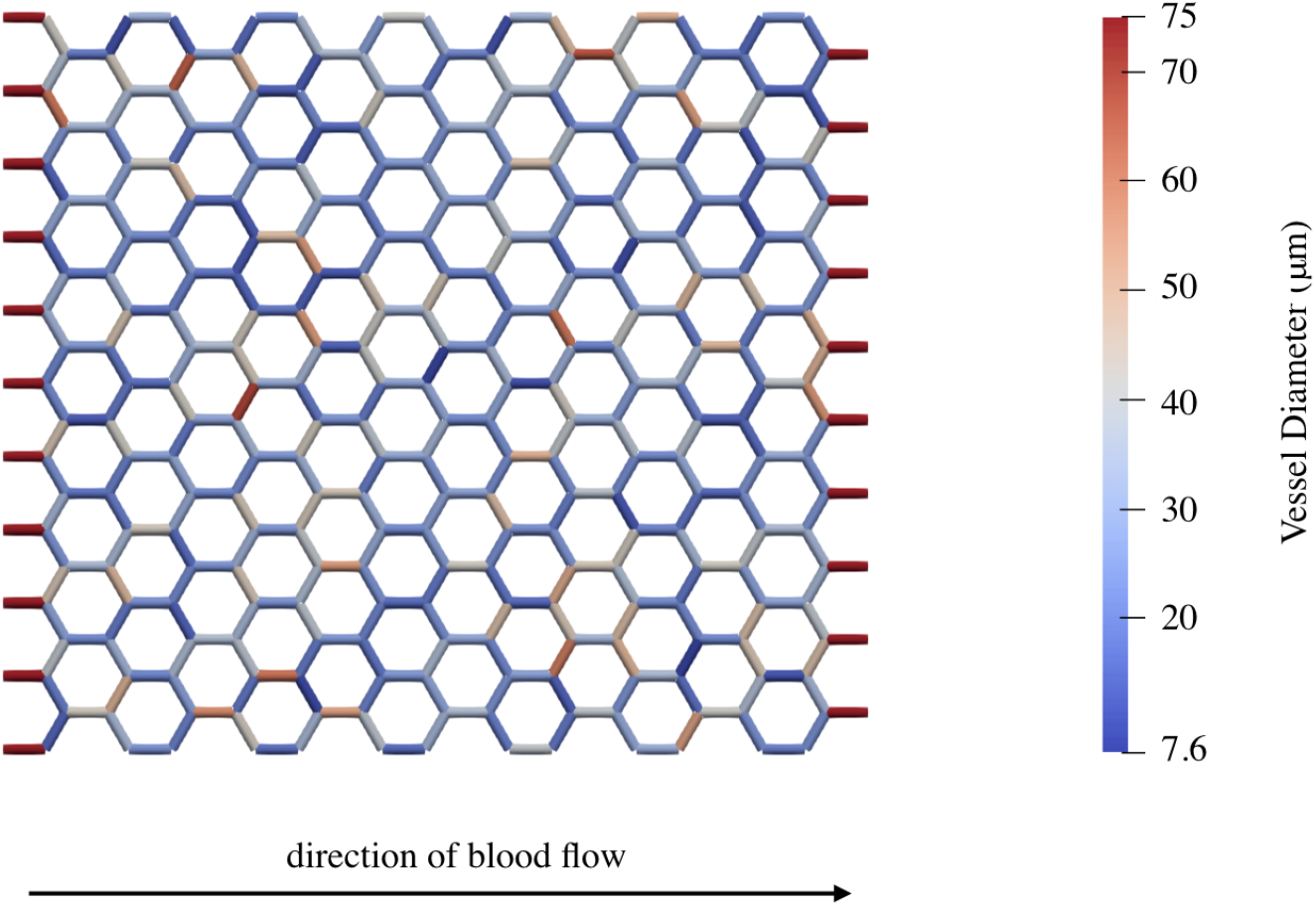
An example of a hexagonal network with diameters taken from a log-normal distribution. Blood flows in from the leftmost nodes and out from the rightmost.

As observed in biological tumours, the distribution of vessel diameters resembles a log-normal distribution [13, 32]. Therefore, vessel diameters in our synthetic networks are randomly sampled from a log-normal distribution with the probability density function:

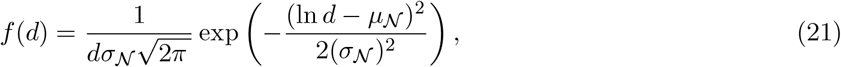

where *d* is the vessel diameter,

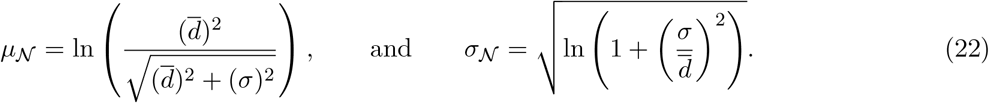

We vary the mean 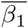 and standard deviation (*σ*) of the diameter distribution to cover the ranges exhibited by the biological tumours (Figure 20). We also impose a minimum and maximum limit for the distribution. The minimum is set to 7.39 μm (the lowest vessel diameter observed in the experiments) and the maximum to 75 μm (the inlet diameter of the forking network).

**Figure 20:**
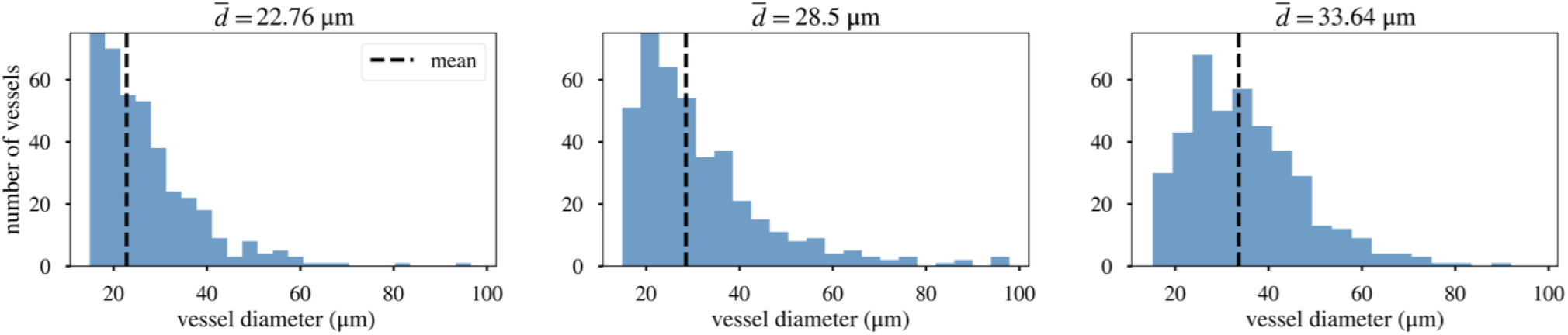
Examples of diameter distributions with a consistent standard deviation (*σ* = 13.23 μm) centred on varying mean diameters for hexagonal networks.

The inlet and outlet vessels are always assigned a diameter of 75 μm in line with the forking network to prevent them from being pruned until no other vessels remain. Results for each network configuration are averaged across 100 randomly sampled distributions.

## D The effect of heterogeneity and hierarchy

Although we lack the experimental data to investigate the effects of vascular diameter heterogeneity and hierarchy in biological networks, we have investigated these features in our synthetic networks. We discuss our results below.

### D.1 Heterogeneity results in increased and persistent perfusion

As mentioned in Section 3.1, diameter heterogeneity in forking networks is modulated by making one daughter vessel *α*-times thicker than the other. We find that as *α* increases, the relative improvement in perfusion fraction (Δ_%_𝒫) also increases (Figure 10). Unlike our analysis of mean diameter 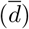, however, the initial perfusion fractions (𝒫_0_) of networks for any given 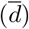 are similar (with the exception of 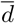 = 33.65 μm) across different values of *α* (Figure 12). Therefore, the initial perfusion fraction does not explain why higher levels of heterogeneity lead to a greater relative improvement.

Examining the network composition helps explain this observation (Figure 10). The maximum perfusion fraction attained over pruning 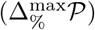 represents the point at which the ratio between the number of perfused vessels and the total number of vessels is greatest (see Equation (1)). Since the latter encapsulates the former, this point also represents the stage at which the ratio between the number of perfused and hypoperfused vessels is greatest. As *α* increases, the hypoperfused vessel count attains smaller minimum values and, therefore, the 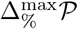 reaches higher values. These successively smaller minima are reached because more heterogeneous networks feature greater proportions of small vessels, which carry blood at a flow rate below the perfusion threshold (Figure 16). Therefore, there exist a greater number of hypoperfused vessels that can be pruned to raise Δ_%_ before pruning perfused vessels.

We also note that networks with smaller values of *α* require fewer vessels to be pruned before zero perfusion is reached. The network exhibits zero perfusion when there is no connected path between the inlet and outlet vessels. As the diameter asymmetry induced by *α* increases, several vessels in the middle of the network become thicker than vessels in other generations and remain unpruned for longer. Therefore, a path between the inlet and outlet is preserved through later stages of pruning in more heterogeneous forking networks.

### D.2 Hierarchy is conducive to perfusion enhancement

The final architectural feature to be evaluated is network hierarchy. Our forking network exhibits diameter hierarchy in the form of a branching architecture, in which parent vessels are thicker (and longer) than their daughter vessels. This order is absent in hexagonal networks (Figure 19).

We observed less pronounced increases in perfusion for non-hierarchical networks than for hierarchical networks (Figure 21). Inspection of the network compositions reveals that the minor increases in perfusion (Δ_%_*𝒫*) are due to the removal of hypoperfused vessels rather than the rerouting of flow. As in the forking networks, 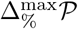is larger in networks with lower mean diameters.

**Figure 21:**
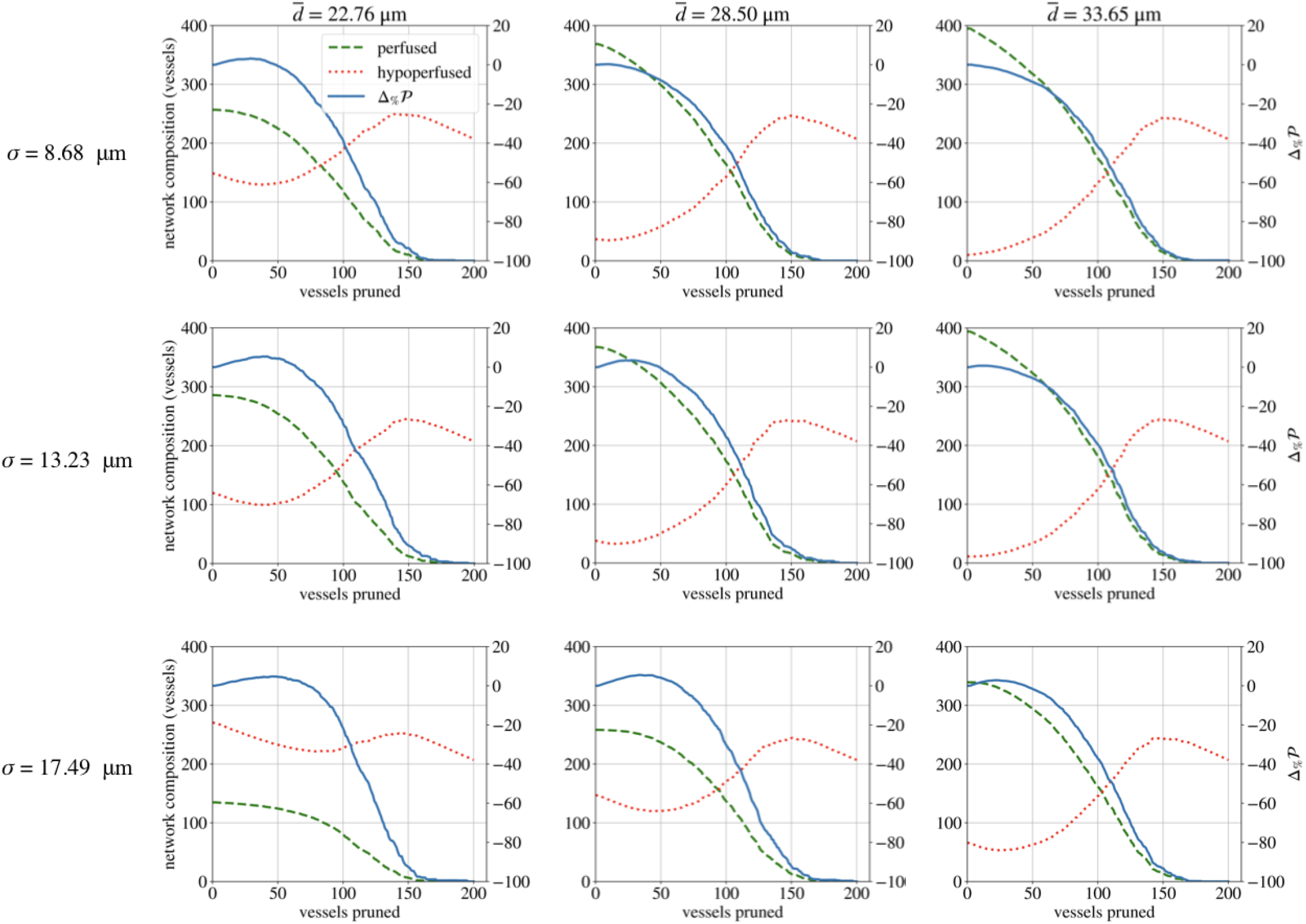
The perfusion response (Δ_%_*𝒫*) of the hexagonal network is largely monotonic during pruning, regardless of the standard deviation of the initial diameter distribution (*σ*).

## E Model parameters

Parameters common to the forking and hexagonal simulations can be found in Table 6. Parameters specific to the forking network can be found in Table 7, while the specifications for the hexagonal network are documented in Table 8.

**Table 6:** Parameter values common to both forking and hexagonal simulations.

| Parameter | Description | Value | Unit | Reference |
| --- | --- | --- | --- | --- |
| $\bar{d}$ | mean network diameter | 22.76, 28.50, 33.65 | $\mu\text{m}$ | Appendix B.1 |
| $d_{inlet}$ | inlet diameter | 75 | $\mu\text{m}$ | Appendix B.1 |
| $\mu_p$ | plasma viscosity | $10^{-3}$ | $\text{kg m}^{-1}\text{s}^{-1}$ | [13] |
| $H_{inlet}$ | inlet haematocrit | 0.45 | - | [13] |
| $p_{inlet}$ | inlet pressure | 3333 | Pa | [57] |
| $p_{outlet}$ | outlet pressure | 2000 | Pa | [57] |

**Table 7:** Parameter values for the forking network architecture.

| Parameter | Description | Value | Unit | Reference |
| --- | --- | --- | --- | --- |
| $\lambda$ | ratio of vessel length to diameter | 4 | - | [13] |
| $\alpha$ | relative thickness of daughter vessels | 1.1, 1.2, 1.3 | - | Appendix B.1 |
| $Q_{min}$ | perfusion threshold | $3 \times 10^{-12}$ | $\text{m}^3 \text{s}^{-1}$ | Section 3.3 |

**Table 8:** Parameter values for the hexagonal network architecture.

| Parameter | Description | Value | Unit | Reference |
| --- | --- | --- | --- | --- |
| $\sigma$ | SD of diameter distribution | 8.68, 13.23, 17.49 | $\mu\text{m}$ | Appendix C |
| $d_{min}$ | minimum diameter | 7.39 | $\mu\text{m}$ | Appendix C |
| $d_{max}$ | maximum diameter | 75 | $\mu\text{m}$ | Appendix C |
| $L_{hex}$ | vessel length | 100 | $\mu\text{m}$ | Appendix C |
| $Q_{min}$ | perfusion threshold | $3 \times 10^{-13}$ | $\text{m}^3 \text{s}^{-1}$ | Section 3.3 |

